# Conjugation of *cis*-OPDA with amino acids is a conserved pathway affecting *cis*-OPDA homeostasis upon stress responses

**DOI:** 10.1101/2023.07.18.549545

**Authors:** Federica Brunoni, Jitka Široká, Václav Mik, Tomáš Pospíšil, Michaela Kralová, Anita Ament, Markéta Pernisová, Michal Karady, Mohamed Htitich, Minoru Ueda, Kristýna Floková, Claus Wasternack, Miroslav Strnad, Ondřej Novák

## Abstract

Jasmonates (JAs) are a family of oxylipin phytohormones regulating plant development and growth and mediating ‘defense *versus* growth’ responses. The upstream JA biosynthetic precursor *cis*-(+)-12-oxo-phytodienoic acid (*cis*-OPDA) has been reported to act independently of the COI1-mediated JA signaling in several stress-induced and developmental processes. However, its means of perception and metabolism are only partially understood. Furthermore, *cis*-OPDA, but not JA, occurs in non-vascular plant species, such as bryophytes, exhibiting specific functions in defense and development. A few years ago, a low abundant isoleucine analog of the biologically active JA-Ile, OPDA-Ile, was detected in wounded leaves of flowering plants, opening up to the possibility that conjugation of *cis*-OPDA to amino acids might be a relevant mechanism for *cis*-OPDA regulation. Here, we extended the analysis of amino acid conjugates of *cis*-OPDA and identified naturally occurring OPDA-Val, OPDA-Phe, OPDA-Ala, OPDA-Glu, and OPDA-Asp in response to biotic and abiotic stress in Arabidopsis. The newly identified OPDA-amino acid conjugates show *cis*-OPDA-related plant responses in a JAR1-dependent manner. We also discovered that the synthesis and hydrolysis of *cis*-OPDA amino acid conjugates are regulated by members of the amidosynthetase GH3 and the amidohydrolase ILR1/ILL families. Finally, we found that the *cis*-OPDA conjugative pathway already functions in non-vascular plants and gymnosperms. Thus, one level of regulation by which plants modulate *cis*-OPDA homeostasis is the synthesis and hydrolysis of OPDA-amino acid conjugates, which temporarily store *cis*-OPDA in stress responses.

## Introduction

Plant hormones are a structurally unrelated collection of small molecules derived from various essential metabolic pathways. Collectively these compounds control every aspect of plant life, from pattern formation during development to responses to biotic and abiotic stress throughout all plant kingdom (Brunoni et al., 2019a; Blázquez et al., 2020). In the pantheon of plant hormones, jasmonates (JAs) are a family of oxylipin phytohormones produced by oxidative metabolism of polyunsaturated fatty acids, regulating aspects of plant development and growth, such as seed germination, root growth, flowering time, stamen development, and senescence (Wasternack and Hause, 2013; Wasternack and Feussner, 2018). JAs are also produced markedly upon abiotic and biotic stresses, such as wounding, insect herbivory, and pathogen infection (Wasternack and Feussner, 2018). The biosynthesis of JAs is a complex process mediated by sequential enzymatic reactions involving three subcellular compartments. Initially, the biosynthesis process occurs in the plastid, where oxidation of the unsaturated fatty acid α-linolenic acid (18:3) mediated by the enzyme 13-lipoxygenase (LOX) takes place (Wasternack and Feussner, 2018). Subsequent dehydration and cyclization reactions catalyzed by allene oxide synthase (AOS) and allene oxide cyclase (AOC), respectively, form *cis*-(+)-12-oxo-phytodienoic acid (*cis*-OPDA). A parallel biosynthetic pathway, starting with hexadecatrienoic acid (16:3) and undergoing similar reactions, produces dinor-OPDA (dnOPDA; Weber et al., 1997). *cis*-OPDA and dnOPDA are then imported to the peroxisome, where their cyclopentenone ring is reduced by OPDA reductase 3 (OPR3), followed by three cycles of β-oxidation leading to jasmonic acid (JA) formation (Breithaupt et al., 2006). Alternatively, JA formation can occur by conversion of *cis*-OPDA to dnOPDA that in turn undergoes peroxisomal β-oxidation, followed by OPR2-mediated cytosolic reduction, thus bypassing the OPR3 pathway (Chini et al., 2018). In all instances, once JA is formed, it is then conjugated in the cytosol with isoleucine to (+)-7-*iso*-jasmonyl-L-isoleucine (JA-Ile) or with other amino acids by the two members of the acyl acid amide synthetases belonging to the GRETCHEN HAGEN 3 (GH3) family, JASMONATE RESISTANT1 (JAR1/GH3.11) and GH3.10 (Staswick and Tiryaki, 2004; Delfin et al., 2022). Upon stress or developmentally regulated processes, JA-Ile level increases, thus leading to the formation of the jasmonate co-receptor complex formed by the F-box coronatine insensitive 1 (COI1) and the jasmonate ZIM domain (JAZ) in which JA-Ile acts as ‘molecular glue’ (Fonseca et al., 2009; Yan et al., 2009). JA-Ile-mediated COI1-JAZ interaction triggers ubiquitination of JAZ repressors and their degradation by the proteasome, thus activating the transcription factors that regulate the JA-specific physiological responses (Thines et al., 2007; Chini et al., 2007; Monte et al., 2022). JA-Ile is turned over by two inducible and intertwined catabolic pathways. One is oxidative and mediated by cytochrome P450 enzymes of the subfamily 94 (CYP94), and the other one proceeds via deconjugation by the three members of the amidohydrolase indole-3-acetic acid (IAA)-LEUCINE RESISTANT 1 (ILR1) and ILR1-like (ILL) family, ILL6, IAR3 and ILR1 (Widemann et al., 2013; Zhang et al., 2016). An additional catabolic pathway based on JA oxidation to OH-JA, catalyzed by members of the 2-oxoglutarate/Fe(II) dioxygenase family, regulates JA turnover upstream of the JA-Ile formation (Caarls et al., 2017).

Although *cis*-OPDA functions primarily as a JA precursor, it also possesses a signaling role distinct from JA-Ile. Any interaction of *cis*-OPDA via the COI1-JAZ co-receptor complex was excluded, thus supporting the OPDA perception via an alternative route (namely COI1-independent pathway; Fonseca et al., 2009; Yan et al., 2009). Remarkably, in contrast to vascular plants, *Marchantia polymorpha* and *Physcomitrium patens* produce *cis*-OPDA and dnOPDA, but not JA and JA-Ile, and isomeric forms of dnOPDA were identified as ligands of MpCOI1 (Monte et al., 2018; Kneeshaw et al., 2022; Široká et al., 2022). Several physiological processes, such as tendril coiling of *Bryonia dioica*, seed germination, thermotolerance, stomatal opening, and response to pathogens, have been attributed to *cis*-OPDA (Weiler et al., 1993; Dave et al., 2011, 2016; Monte et al., 2020; Chang et al., 2023). Uncoupling OPDA and JA-Ile synthesis was proven to be particularly challenging in *Arabidopsis thaliana*. Many published studies of Arabidopsis oxylipin signaling rely on the *opr3-1* mutant, later discovered to be a conditional allele not completely impaired in JA-Ile biosynthesis (Chehab et al., 2011). A knockout *opr3* sterile mutant (*opr3-3*) was subsequently identified, and fertility was restored upon JA treatment (Chini et al., 2018). It has been proposed that *cis*-OPDA exerts its regulatory function by reversibly binding the cyclophilin 20-3 (CYP20-3), thus stabilizing enzymes involved in cysteine synthesis (Park et al., 2013). This event triggers the glutathione (GSH) level increase, thus determining redox changes in the plastid and cytosol and *cis*-OPDA-mediated TGA transcription factor activation (Jimenez-Aleman et al., 2022). Several conjugates of *cis*-OPDA, such as OPDA-GSH and OPDA-amino acid conjugates (namely OPDA-aa), have been identified (Ohkama-Ohtsu et al., 2011; Floková et al., 2016; Shinya et al., 2022). On the one hand, conjugation of *cis*-OPDA with GSH is described as a possible detoxification mechanism of *cis*-OPDA after stress, as this conjugate accumulates in the vacuole, and its reactivity is linked to cellular redox homeostasis (Ohkama-Ohtsu et al., 2011; Park et al., 2013). On the other hand, since conjugation of JA with amino acids is required for its bioactivity and homeostasis, it is tempting to speculate that a similar scenario may occur in the case of *cis*-OPDA-specific response. In Arabidopsis, OPDA-Ile was identified as a low abundant metabolite in wounded leaves and proposed to act as a regulatory molecule in a JA-independent manner (Floková et al., 2016; Arnold et al., 2016). OPDA-Ile was shown to induce the expression of genes encoding for the C_2_H_2_-type zinc finger transcription factor, ZAT10, and the glutaredoxin, GRX480, that were previously identified as *cis*-OPDA-inducible (Taki et al., 2005; Park et al., 2013; Arnold et al., 2016). Several OPDA-aa have recently been putatively identified in rice and proposed as non-canonical signaling molecules for producing phytoalexins in coordination with innate chitin signaling (Shinya et al., 2022). Overall, these studies corroborate the finding that conjugation of *cis*-OPDA occurs in plants in response to different stimuli; although the mechanism by which OPDA-aa are formed, their role and function in plants remain elusive and poorly understood.

Here, we extended the analysis of amino acid conjugates of *cis*-OPDA and described naturally occurring OPDA-Val, OPDA-Phe, OPDA-Ala, OPDA-Glu, and OPDA-Asp. These conjugates accumulated upon wounding stress, *cis*-OPDA homeostasis perturbation, and fungal pathogen infection in Arabidopsis. Like free *cis*-OPDA, the newly identified OPDA-aa show JAR1-dependent growth-inhibitory effect, trigger the JAZ repressor degradation and influence the expression of a subset of OPDA- and JA-responsive genes. Moreover, we showed that members of the GH3 and ILR1/ILL families catalyze the conjugation of *cis*-OPDA to amino acids and hydrolysis of the OPDA-aa, respectively. We also demonstrated that the conjugation of *cis*-OPDA with amino acid is a metabolic pathway that occurred early during land plant evolution. It already operates in the moss *P. patens*, and the gymnosperm *Picea abies*. Thus, similarly to other phytohormones, one level of regulation by which plants modulate *cis*-OPDA homeostasis appears to be the synthesis and hydrolysis of OPDA-aa, which function in the temporary storage of *cis*-OPDA in stress responses.

## Results

### OPDA-aa accumulate in response to biotic and abiotic stress in Arabidopsis

To explore whether the occurrence of OPDA-aa is a specific or a broad-spectrum metabolic response to stressful events in Arabidopsis, we monitored the formation of OPDA-aa in several well-studied conditions known to alter JA and *cis*-OPDA homeostasis, including leaf wounding, pathogen infection, and chemical treatment. We previously demonstrated that OPDA-Asp, OPDA-Glu, OPDA-Ile, OPDA-Phe, and OPDA-Val were detected in Arabidopsis wounded leaves (Mik et al., 2023). These OPDA-aa did not accumulate in the earliest time points post-wounding but were instead found in the latest time points after injury, suggesting that OPDA-aa formation might follow a kinetics that differs from the rapid JA-Ile wounding response (Mik et al., 2023; Koo et al., 2009). Therefore, we performed a new experiment and focused on OPDA-aa occurrence in the time points ranging from 30 min to 4 h after wounding. We found *cis*-OPDA conjugated with Ala, Glu, and Phe, while OPDA-Ile, OPDA-Val, OPDA-Asp, and OPDA-Trp were not detected in wounded wild-type Arabidopsis leaves under our experimental conditions (**Figure 1A**). Except for OPDA-Ala, which accumulated 30 min after wounding and kept increasing after 4 h, OPDA-Glu, and OPDA-Phe were detected by 2 h and peaked by 4 h post-wounding. We also monitored the amino acid conjugation of JA. We observed that, upon wounding, OPDA-aa accumulated with kinetics that was more similar to the less abundant JA-Ala and JA-Val than the typical rapid JA-Ile accumulation (**Supplemental Figure S1A**; Koo et al., 2009). OPDA-aa levels ranged between 1 to 10 pmol/g fresh weight (FW), consistent with previous results (Mik et al., 2023). Some differences were also observed (**Figure 1A**; Mik et al., 2023). OPDA-Ile, OPDA-Val, and OPDA-Asp were not detected, and OPDA-Ala was found upon wounding in this study, while OPDA-Ile, OPDA-Val, and OPDA-Asp occurred and OPDA-Ala was not identified in the previous study. This could be explained by the different experimental settings adopted for the wounding experiment; first of all, older plants grown under a 12/12-h photoperiod were used in this study whereas younger plants grown under long photoperiod (16/8 h) were used in the previous study. Secondly, mechanical stress was performed on expanded leaves by wounding three times the midvein in this study, while leaves were wounded once on the one side of the central vein in the previous experiment. To assess whether the formation of OPDA-aa could be elicited upon fungal attack, we inoculated Arabidopsis plantlets with spores of *Botrytis cinerea*, a broad-spectrum pathogen widely used to study plants response to biotic stress (Chini et al., 2018). We found *cis*-OPDA conjugated to Ala, Asp, Glu, and Ile only in *B. cinerea*-inoculated plants, while none of the OPDA-aa was detected in mock-treated plants (**Figure 1B**). OPDA-Phe, OPDA-Val, and OPDA-Trp were not detected in response to this pathogen or mock samples. We also quantified JA-aa and observed high levels of JA-Ala and JA-Glu, and that accumulation JA-Ile, JA-Asp, JA-Gly, and JA-Val occurred to a lesser extent (**Supplemental Figure S1B**). On the contrary, a similar range of OPDA-aa levels was recorded by wounding and fungal infection (**Figure 1, A and B**). A wider variety of OPDA-aa was detected after exogenous treatment with *cis*-(±)-OPDA, compared to the response observed in wounded- and infected-plants (**Figure 1**). All the inspected OPDA-aa, except OPDA-Trp, were rapidly formed upon application of *cis*-(±)-OPDA 5 min after treatment, with OPDA-Asp and OPDA-Glu being the most abundant OPDA-aa (**Figure 1C**). The range of accumulated OPDA-aa levels was much less than that of JA-Ile (**Figure 1A**; Koo et al., 2009). Nevertheless, these results indicate that *cis*-OPDA amino acid conjugation is a metabolic mechanism plants adopt to respond to a wide range of physiological conditions perturbing the *cis*-OPDA and JA homeostasis.

**Figure 1.**
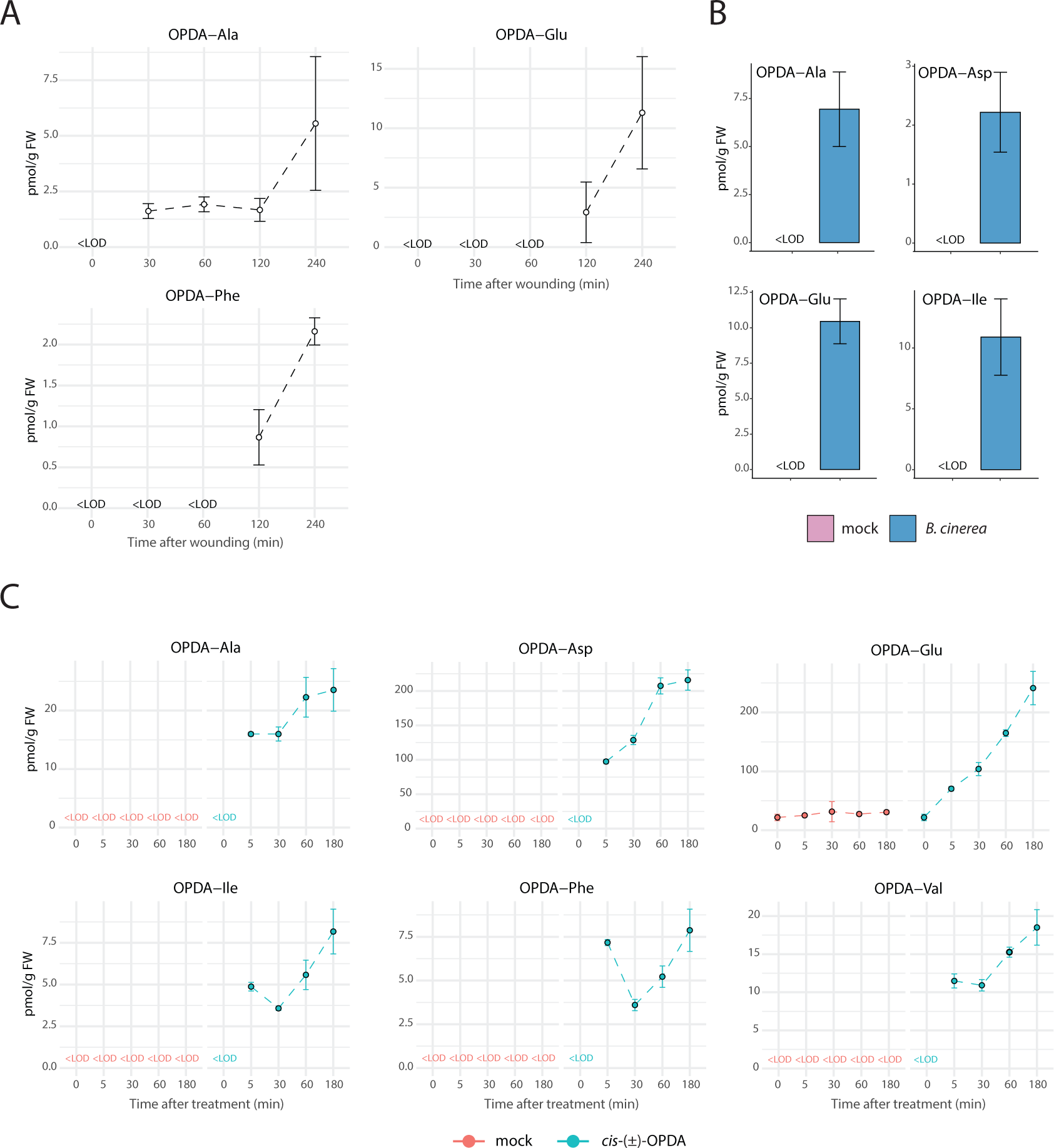
OPDA-aa accumulate in Arabidopsis plants during stress responses. A, Time-course accumulation of indicated OPDA-aa in Col-0 after leaf wounding. Six-week-old plants were wounded, and damaged leaves were collected after the indicated times. B, Accumulation of indicated OPDA-aa in Col-0 plants infected with *B. cinerea* 3 days after inoculation. C, Time-course accumulation of indicated OPDA-aa after exogenous treatment with 50 µM *cis*-(±)-OPDA. Plants were sampled after the indicated times. OPDA-aa levels are expressed as pmoles per gram of plant fresh weight (FW). Mean ± SD (*n*=3). Below the limit of detection, <LOD.

### Study of the activity of OPDA-aa in cis-OPDA- and JA-regulated responses

JA and *cis*-OPDA inhibit root growth (Monte et al., 2018). Therefore, a primary root-growth inhibition assay was carried out to examine possible *cis*-OPDA-related phenotype upon treatment with (±)-OPDA-aa. Since OPDA-Trp was not detected in any of the experiments described above (**Figure 1**), this molecule was excluded in the following experiments. Overall root growth was significantly reduced on (±)-OPDA-aa-containing medium compared to mock-treated seedlings. Nonetheless, we observed that treatment with (±)-OPDA-Val, (±)-OPDA-Phe, and (±)-OPDA-Ala impaired root length similarly to *cis*-(±)-OPDA and significantly more than (±)-OPDA-Ile, (±)-OPDA-Asp, and (±)-OPDA-Glu (**Figure 2, A and B**). It should be noted that a racemic mixture of (±)-OPDA-aa, made from racemic *cis*-(±)-OPDA, was used when applied exogenously; thus only 50% of the applied chemicals have the proper stereochemistry and is biologically active. To investigate whether the root-growth inhibitory effect of these (±)-OPDA-aa required *in planta* conjugation of JA to Ile, we also grew seeds of *jar1-11* mutant in the presence of (±)-OPDA-aa. We chose *jar1-11* rather than other JA biosynthetic mutants, such as *opr3-3*, because although both mutants are impaired in the JA-Ile synthesis, *jar1-11* is fertile, whereas *opr3-3* is sterile and thus difficult to work with (Chini et al., 2018). All the (±)-OPDA-aa showed an effect similar to *cis*-(±)-OPDA and (±)-JA, thus partially inhibiting the growth of *jar1-11* mutant, demonstrating that the (±)-OPDA-aa-induced root-growth inhibitory effect requires JA-Ile formation (**Figure 2, A and B**). To test whether these molecules could impact the JA signaling pathway, we studied the effect of (±)-OPDA-aa on hormone-induced degradation of JAZ repressors, a typical COI1-mediated response, using the Arabidopsis *35S:JAZ1*-β-glucuronidase (GUS) and *35S:JAZ9*-GUS reporter lines (Thines et al., 2007). All the (±)-OPDA-aa triggered JAZ1 and JAZ9 protein degradation, similarly to (±)-JA and *cis*-(±)-OPDA, showing that their effect is not restricted to a particular JAZ (**Figure 2C**). We further monitored the effect of (±)-OPDA-aa treatment on expression of genes known to be *cis*-OPDA and JA-responsive markers, such as *thionin* (*THI2.1*), *VEGETATIVE STORAGE PROTEIN1* (*VSP1*), *ZAT10*, *GRX480*, *JAZ5* and *PLANT DEFENSIN1.2* (*PDF1.2*; Staswick and Tiryaki, 2004; Arnold et al., 2016; Chini et al., 2018), and compared their expression level upon treatment with (±)-JA, *cis*-(±)-OPDA or mock (**Figure 3A**). None of the (±)-OPDA-aa triggered the expression of *THI2.1* and *VSP1*, whereas the transcripts of the other marker genes increased, compared with the mock treatment, although with some differences. On the one hand, (±)-OPDA-Val, (±)-OPDA-Phe, and (±)-OPDA-Ala were found equally effective in upregulating the expression of *ZAT10*, *GRX480*, and *PDF1.2* genes, with the induction of the *ZAT10* and *GRX480* expression comparable to *cis*-(±)-OPDA and (±)-JA. On the other hand, no significant upregulation of *PDF1.2* and *JAZ5* expression levels was observed upon (±)-OPDA-Ile, (±)-OPDA-Asp, and (±)-OPDA-Glu, and in general, the response exerted by this group of (±)-OPDA-aa was found consistent (**Figure 3A**). Results from the root-growth inhibition assay and gene expression analysis suggested that the six newly identified OPDA-aa could be divided into two groups (**Figure 2, A and B**; **Figure 3A**). Therefore, we selected OPDA-Val and OPDA-Glu, as representatives of each group to design standards isotopically labeled on the amino acid moiety. These standards were used in a feeding experiment to assess whether plants can hydrolyze these conjugates. Both isotopically labeled (±)-OPDA-aa were taken up by Arabidopsis seedlings, as *cis*-(±)-OPDA-[^13^C_5_,^15^N]*L*-Glu and *cis*-(±)-OPDA-[^13^C_5_,^15^N]*L*-Val accumulated in plant samples, whereas their level decreased in the liquid medium progressively (**Figure 3B**). Free [^13^C_5_,^15^N]*L*-Val was detected 30 min after treatment in Arabidopsis seedlings, indicating that (±)-OPDA-Val is hydrolyzed *in planta*. Conversely, [^13^C_5_,^15^N]*L*-Glu was not detected in plant samples at any time point after treatment. This might suggest that the hydrolytic cleavage does not occur at the amide bond position or that free [^13^C_5_,^15^N]*L*-Glu is either converted to γ-aminobutyric acid or serves as a known immediate major donor for the synthesis of other amino acids and nitrogen-containing compounds in plants (Forde et al., 2007; Kan et al., 2017). Together these results hint that (±)-OPDA-aa exhibit *cis*-OPDA-like activity in a JAR1-dependent manner, thus likely exerting their function upon hydrolysis of the amino acid moiety.

**Figure 2.**
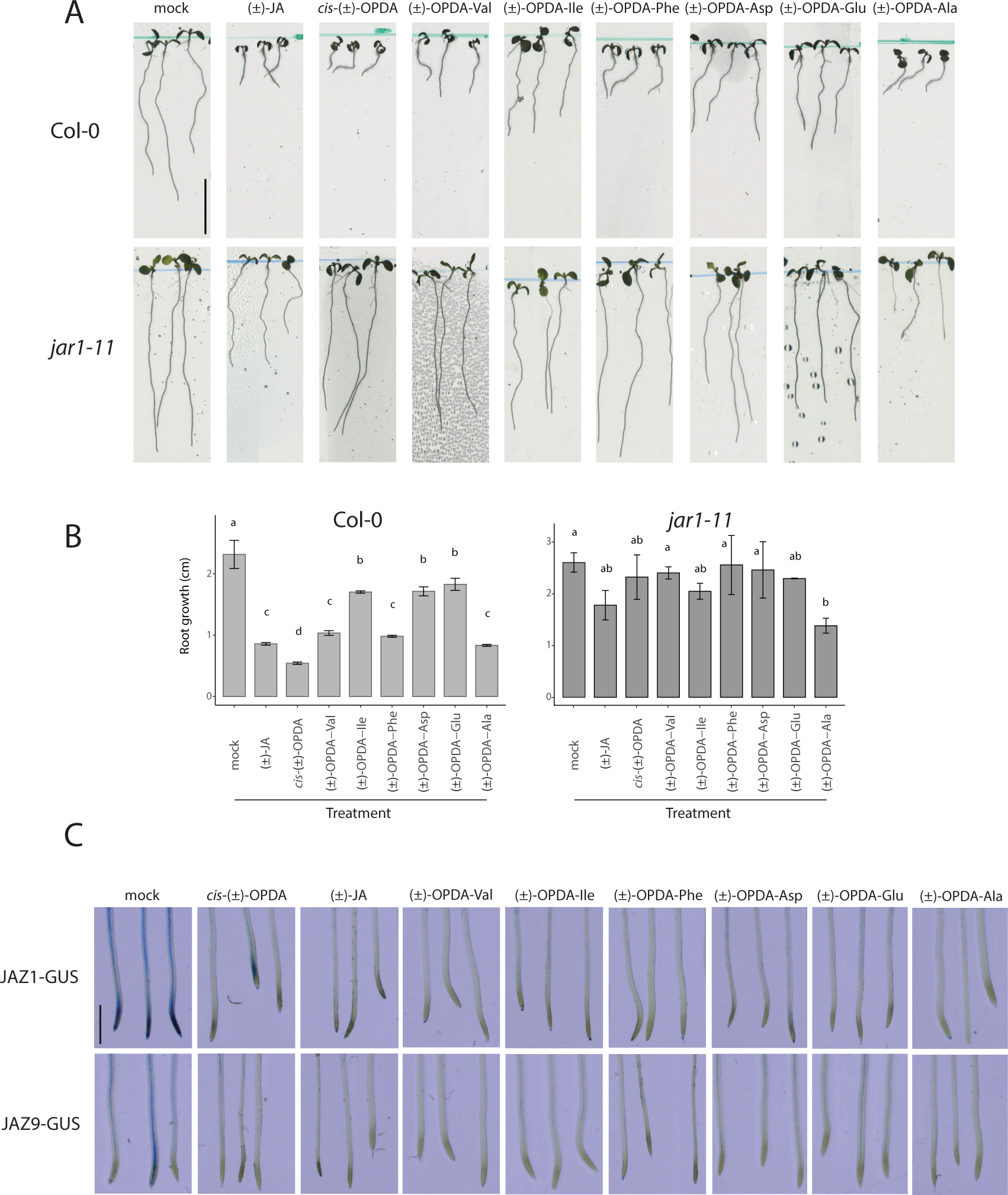
(±)-OPDA-aa exhibit *cis*-OPDA-like activity in a JAR1-dependent manner. A, Wild-type (Col-0) and *jar1-11* mutant seedlings grown on vertical plates in the absence (mock) or presence of 50 µM (±)-JA or 5 µM *cis*-(±)-OPDA, and indicated (±)-OPDA-aa. Scale bar presents 1 cm. B, Root length was measured 7 days after germination. Data are shown as mean ± SD of three biological replicates. Letters indicate significant differences, evaluated by one-way ANOVA/Tukey HSD *post hoc* test (*P*<0.05). C, Representative seedlings of the *35S:JAZ1-GUS* and *35S:JAZ9-GUS* lines were treated with or without 10 µM (±)-JA, *cis*-(±)-OPDA, and indicated (±)-OPDA-aa for 2 h. Scale bar represents 1 mm.

**Figure 3.**
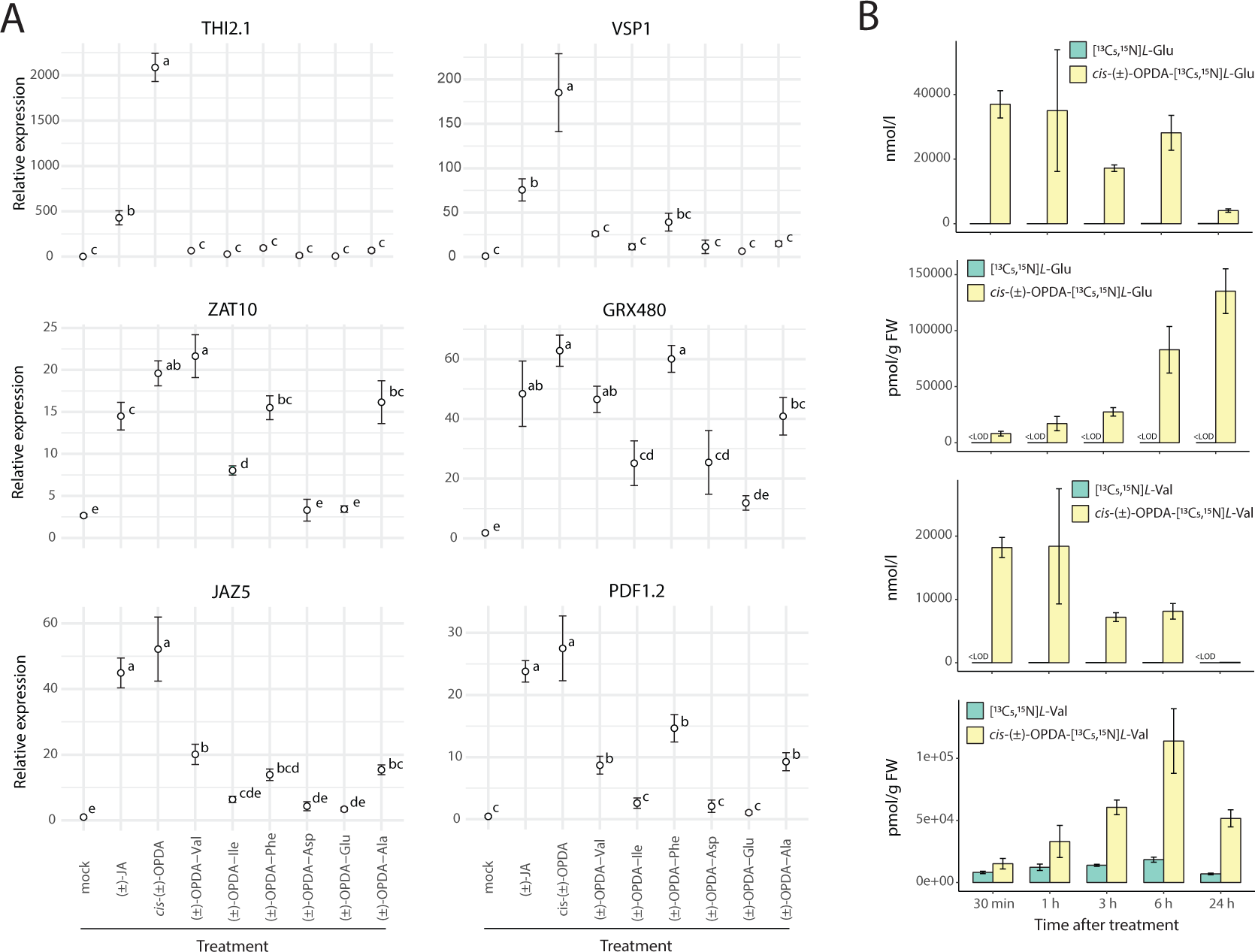
Gene expression analysis of *cis*-OPDA- and JA-marker genes and labeled (±)-OPDA-aa feeding in Arabidopsis. A, Expression of *THI2.1*, *VSP1*, *ZAT10*, *GRX480*, *JAZ5*, and *PDF1.2* after treatment with or without 50 µM (±)-JA, *cis*-(±)-OPDA, and indicated (±)-OPDA-aa in Col-0. Gene expression was measured by RT-qPCR. Data are expressed as relative fold change normalized by *ACT2*. Letters indicate significant differences, evaluated by one-way ANOVA/Tukey HSD *post hoc* test (*P*<0.05). B, Time-course accumulation of *cis*-(±)-OPDA-[^13^C_5_,^15^N]*L*-glutamate, [^13^C_5_,^15^N]*L*-Glu, *cis*-(±)-OPDA-[^13^C_5_,^15^N]*L*-valine and [^13^C_5_,^15^N]*L*-Val in liquid ½ MS medium (nmol/l) and 7-day-old Col-0 seedlings (pmol/g FW) after feeding with 50 µM *cis*-(±)-OPDA-[^13^C_5_,^15^N]*L*-glutamate and *cis*-(±)-OPDA-[^13^C_5_,^15^N]*L*-valine. Samples were collected at the indicated times. Mean ± SD (*n*=3). Below the limit of detection, <LOD.

### Members of the Groups II and III GH3 family conjugate cis-(±)-OPDA with amino acids in an enzymatic assay and in planta

Arabidopsis possesses 19 different GH3 proteins classified into three groups (I, II, and III) based on substrate specificity and sequence homology (Staswick et al., 2005). To study how *cis*-(±)-OPDA is conjugated, we first tested the enzymatic activity of Arabidopsis Group I GH3s with *cis*-(±)-OPDA, as the GH3.10- and GH3.11-mediated JA conjugation is well-documented (Staswick and Tiryaki, 2004; Delfin et al., 2022). Both recombinant enzymes could conjugate preferentially (±)-JA with Ile, and to a minor extent, with other amino acids (**Supplemental Dataset S1**, **Supplemental Table S1**), confirming published results (Staswick and Tiryaki, 2004; Delfin et al., 2022). Surprisingly, none of these enzymes displayed activity towards *cis*-(±)-OPDA, and only a minimal amount above the limit of detection of (±)-OPDA-Ile was found (**Supplemental Dataset S1**). To investigate whether AtGH3.11 could mediate conjugation of *cis*-OPDA *in planta*, we fed wild-type and *jar1-11* mutant seedlings with (±)-JA and *cis*-(±)-OPDA and inspected the formation of OPDA-aa after 3 h. While the accumulation of JA-aa, such as JA-Val and JA-Ile, was significantly reduced in *jar1-11* mutant upon treatment with (±)-JA compared with wild-type, as expected (Staswick and Tiryaki, 2004; Delfin et al., 2022), OPDA-aa levels were not decreased in the *jar1-11* mutant (**Supplemental Figure S2**), thus confirming that GH3.11 catalyzes the conjugation of JA but not of *cis*-OPDA. We then investigated whether the other members of the GH3 family could conjugate *cis*-(±)-OPDA with amino acids. Among Group II, known to conjugate mainly auxins (Staswick et al., 2005; Brunoni et al., 2023), AtGH3.1, AtGH3.2, AtGH3.3, AtGH3.4, AtGH3.5, and AtGH3.6 displayed the highest activity with *cis*-(±)-OPDA and Asp, and AtGH3.17 conjugated mainly *cis*-(±)-OPDA with Glu (**Figure 4A**; **Supplemental Figure S3A**). Among Group III, AtGH3.12 could conjugate *cis*-(±)-OPDA with Glu primarily and secondly with Val, Ile, and Phe, whereas AtGH3.14 and AtGH3.15 displayed a very similar activity in forming (±)-OPDA-Ala, (±)-OPDA-Trp, and (±)-OPDA-Ile primarily, and to a lesser extent (±)-OPDA-Phe (**Supplemental Figure S3A**). Notably, among the AtGH3s catalyzing the conjugation of *cis*-(±)-OPDA, only those belonging to Group III were found highly active toward (±)-JA (**Supplemental Dataset S1**; **Supplemental Table S1**), supporting previous findings (Westfall et al., 2016; Sherp et al., 2018). To assess whether GH3s conjugate *cis*-OPDA in plants and to overcome gene redundancy interference, we fed wild-type Col-0 and *gh3.1,2,3,4,5,6* (*gh3* sextuple) mutant seedlings with *cis*-(±)-OPDA and followed the formation of OPDA-aa after 3 h. Similar levels of the OPDA-aa were recorded in the wild-type and in the *gh3* sextuple mutant (**Supplemental Figure S2B**), except for OPDA-Asp, whose formation was abolished entirely in the *gh3* sextuple mutant (**Figure 4B**). This data indicates that AtGH3.1, AtGH3.2, AtGH3.3, AtGH3.4, AtGH3.5, and AtGH3.6 are the GH3 enzymes responsible for the conjugation of *cis*-OPDA with Asp in Arabidopsis plants and that the other OPDA-aa (*i.e.*, OPDA-Ala, OPDA-Glu, OPDA-Val, OPDA-Ile and OPDA-Phe) are formed very likely by AtGH3.17 and members of the Group III GH3s.

**Figure 4.**
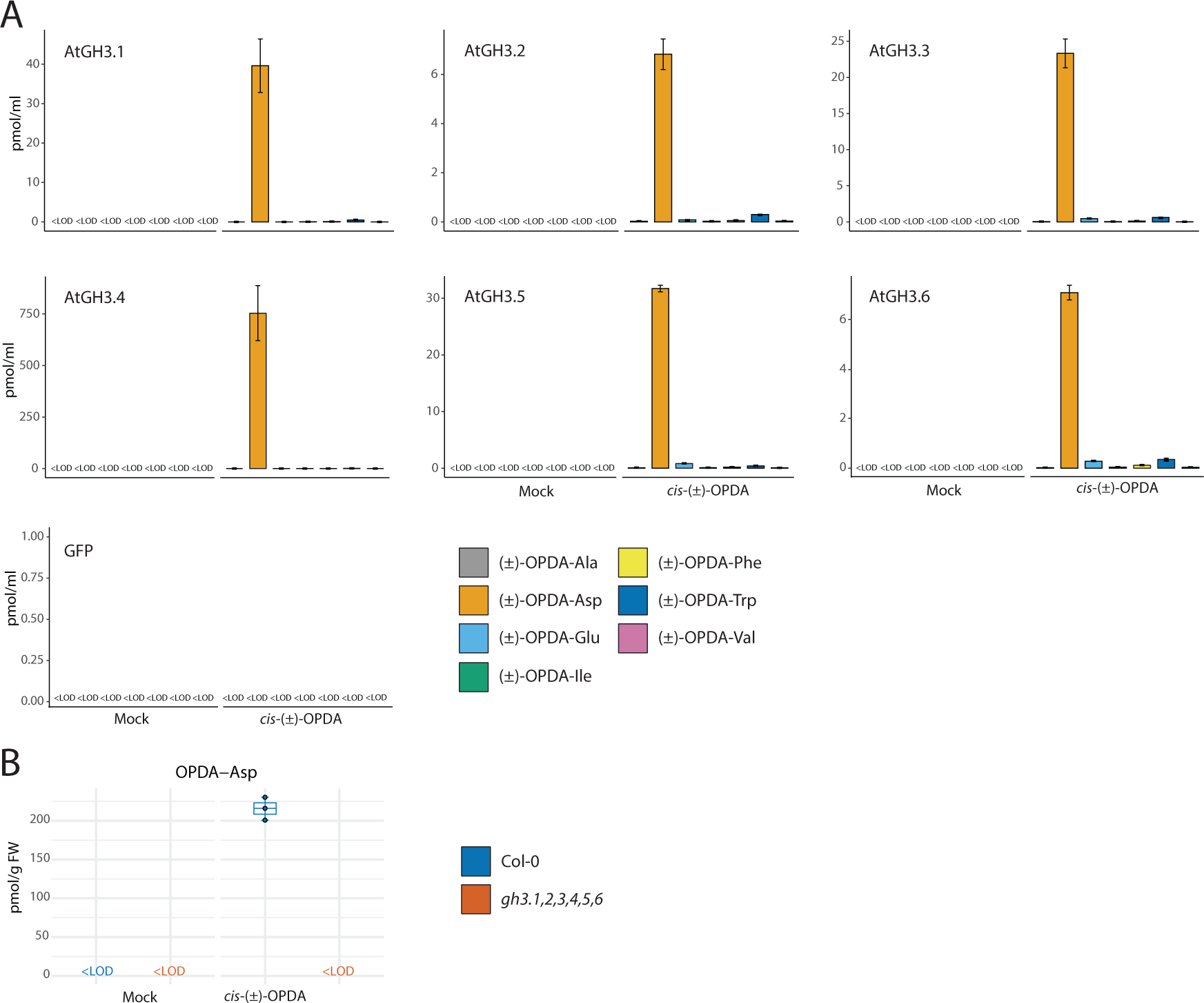
Group II GH3 proteins are responsible for *cis*-(±)-OPDA conjugation with Asp in Arabidopsis. A, Analysis of (±)-OPDA-aa synthesized by recombinant AtGH3.1, AtGH3.2, AtGH3.3, AtGH3.4, AtGH3.5, and AtGH3.6 in the bacterial assay. The cell lysate was incubated with or without 0.1 mM *cis*-(±)-OPDA and GH3 cofactor mixture for 5 h at 30 °C. The bacterial assay carried out with cell lysate from GFP-producing bacteria was used as a negative control. Cell lysate without *cis*-(±)-OPDA and cofactor mixture was used as a mock sample. (±)-OPDA-aa level is expressed as pmol/ml. B, Formation of OPDA-Asp after feeding of seven-day-old Arabidopsis Col-0 and *gh3* sextuple mutant (*gh3.1,gh3.2,gh3.3,gh3.4,gh3.5,gh3.6*) with or without 50 µM *cis*-(±)-OPDA for 3 h. OPDA-Asp level is expressed as pmol/g FW. Horizontal lines in the box plots are medians, boxes show the upper and lower quartiles, and whiskers show the entire data range. Mean ± SD (*n*=3). Below the limit of detection, <LOD.

### Members of the ILR1/ILL family hydrolyze OPDA-aa in an enzymatic assay and in planta upon wounding

Results from the feeding experiment with isotopically labeled (±)-OPDA-aa showed that OPDA-aa can be hydrolyzed *in planta* (**Figure 3B**). A subset of JA-inducible amidohydrolases of the ILR1/ILL family was previously described to catalyze the hydrolysis of JA-aa upon wounding in Arabidopsis leaves (Widemann et al., 2013; Zhang et al., 2016). Therefore, in the following experiments, we investigated the possible involvement of Arabidopsis ILR1, IAR3, ILL2, and ILL6 in the cleavage of OPDA-aa. To study their catalytic activities, recombinant Arabidopsis amidohydrolases were produced in *E. coli*, and conditions for the hydrolysis assay were first determined by incubating the recombinant protein-producing cell lysate with IAA-Ala, confirming previously published results (**Supplemental Dataset S1**; LeClere et al., 2002; Zhang et al., 2016). Next, the activity of these four hydrolases was tested against (±)-OPDA-aa. AtILL2 and AtILR1 hydrolyzed all six (±)-OPDA-aa comparably well (**Figure 5A**). While AtILL2 displayed the highest activity toward (±)-OPDA-Ala, (±)-OPDA-Asp, and (±)-OPDA-Phe, AtILR1 hydrolyzed preferentially (±)-OPDA-Ala, (±)-OPDA-Asp, and (±)-OPDA-Glu. AtIAR3 was able to hydrolyze primarily (±)-OPDA-Ala, (±)-OPDA-Asp, and (±)-OPDA-Val, and to a lesser extent (±)-OPDA-Glu and (±)-OPDA-Phe. AtILL6 showed the highest activity toward (±)-OPDA-Glu, and secondly (±)-OPDA-Phe. Its activity toward (±)-OPDA-Ala and (±)-OPDA-Asp could not be clearly ascribed to AtILL6, as similar *cis*-(±)-OPDA levels were recorded in the GFP-producing bacteria after incubation with (±)-OPDA-Ala and (±)-OPDA–Asp, suggesting the possible presence of an endogenous substrate-associated bacterial machinery (Brunoni et al., 2019b). This enzymatic activity study shows that all the four tested members of the ILR1/ILL family can cleave several OPDA-aa, highlights overlapping but distinct substrate specificities for various amino acid conjugates of *cis*-(±)-OPDA and predicts their possible involvement in *cis*-OPDA homeostasis. To assess whether these hydrolases contributed to regulating the OPDA-aa levels in plants, we carried out a time course OPDA-aa profiling on *ill6-2* single and *ilr1-1,iar3-2,ill2-1* triple knockout mutants upon leaf wounding. OPDA-Ala, OPDA-Glu, OPDA-Ile, and OPDA-Phe accumulated in *ill6-2* mutant already 30 min post-wounding; their level was always higher than Col-0 throughout the time course whereas OPDA-Val was detected only in the *ill6-2* mutant as its level was always below the limit of detection in the Col-0 (**Figure 5B**). Considering the low hydrolytic activity of recombinant AtILL6 against OPDA-Ala and OPDA-Ile, it was surprising that these two OPDA-aa accumulated in the *ill6-2* mutant (**Figure 5**). Nonetheless, a similar discrepancy between *in vitro* and *in vivo* assays was already reported for ILL6 (Zhang et al., 2016). OPDA-Ala and OPDA-Phe were detected in the *ilr1-1,iar3-2,ill2-1* triple mutant 30 min and 1 h after wounding, respectively, their level was steady over time and greater in the triple mutant than Ws. OPDA-Glu was found 4 h after injury only in the triple mutant, and no accumulation in Ws was found. OPDA-Asp was not detected in any of the mutant lines or the wild-types. Moreover, leaf wounding of the *ilr1-1*, *iar3-2*, and *ill2-1* single knockout mutants showed that OPDA-Ala and OPDA-Phe accumulated in all three single mutants. In contrast, OPDA-Glu was detected only in the *iar3-2* and *ill2-1* mutants (**Supplemental Figure S4**). These findings suggest that ILR1, IAR3, ILL2, and ILL6 are all involved in the hydrolysis of several OPDA-aa and that OPDA-aa are rapidly cleaved by these amidohydrolases upon wounding.

**Figure 5.**
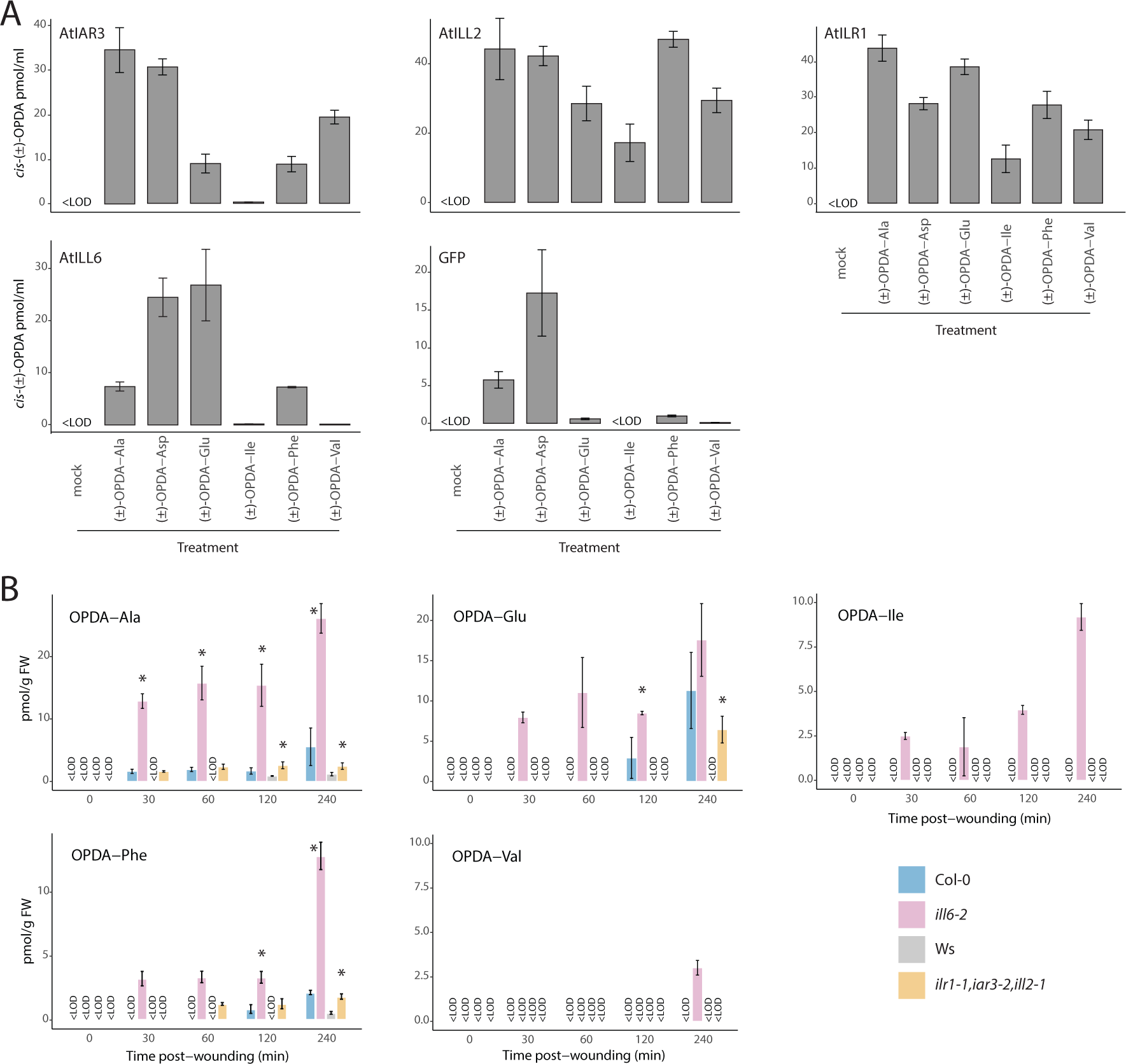
ILR1/ILL enzymes are involved in *cis*-OPDA release from OPDA-aa in Arabidopsis upon wounding. A, Analysis of release of *cis*-(±)-OPDA by recombinant AtIAR3, AtILL2, AtILR1, and AtILL6 in the bacterial assay. The cell lysate was incubated with or without 0.1 mM (±)-OPDA-aa and 1 mM MgCl_2_ for 5 h at 30 °C. The bacterial assay carried out with the cell lysate from GFP-producing bacteria was used as a negative control. Cell lysate without (±)-OPDA-aa and MgCl_2_ was used as a mock sample. *cis*-(±)-OPDA level is expressed as pmol/ml. B, Time-course accumulation of indicated OPDA-aa in wild-types (Col-0, Ws), *ill6-2* single (Col-0 background) and *ilr-1,iar3-2,ill2-1* triple (Ws background) knockout mutants after leaf wounding. Six-week-old plants were wounded, and damaged leaves were collected after the indicated times. Asterisk indicates statistically significant differences, as determined by Student’s *t*-test (Col-0 vs *ill6-2*, Ws vs *ilr1-1,iar3-2,ill2-1*; *P*<0.05). OPDA-aa concentrations are given as pmoles per gram fresh weight (FW). Mean ± SD (*n*=3). Below the limit of detection, <LOD.

### OPDA amino acid conjugation is a conserved mechanism that occurred early during land plant evolution

The occurrence of *cis*-OPDA, but not JA/JA-Ile, is described in different non-vascular species, including bryophytes, and it has been proposed that JA-Ile biosynthesis emerged in lycophytes (Chini et al., 2023). To clarify whether the conjugation of *cis*-OPDA is a conserved metabolic pathway, we inspected the occurrence of this pathway in evolutionarily distant species, such as the moss *P. patens* and the conifer *Picea abies*, as models for bryophytes and gymnosperms, respectively. *P. patens* possesses only two GH3 enzymes (Ludwig-Müller et al., 2009). We first tested the activity of recombinant PpGH3s with *cis*-(±)-OPDA and observed that both isoforms could conjugate *cis*-(±)-OPDA (**Figure 6A**). PpGH3.1 catalyzed the conjugation of *cis*-(±)-OPDA with Ala, whereas PpGH3.2 mainly with Glu and to a much lesser extent with Asp. PpGH3.2, but not PpGH3.1, could also accept (±)-JA as a substrate for conjugation, mainly with Glu (**Supplemental Dataset S1**; **Supplemental Table S1**), supporting previous findings (Ludwig-Müller et al., 2009). A feeding experiment with *cis*-(±)-OPDA of moss wild-type and *gh3* single and double knockout mutant gametophores revealed that OPDA-Glu and OPDA-Asp accumulated in wild-type and the *gh3.1* mutant upon *cis*-(±)-OPDA treatment. In contrast, the level of OPDA-Glu was deficient and no OPDA-Asp was detected in the *gh3.2* single and *gh3* double mutants, confirming that PpGH3.2, but not PpGH3.1, catalyzes the conjugation of *cis*-OPDA with Glu and Asp (**Figure 6B**). Isomerization of *cis*-OPDA to 12-oxo-9(13),15(Z)-phytodienoic acid (*iso*-OPDA) was very recently reported to occur in *P. patens* with *iso*-OPDA described as a predominant oxylipin in this species (Mukhtarova et al., 2020). Thus, we monitored the formation of *iso*-OPDA after treatment of moss gametophores with *cis*-(±)-OPDA and confirmed the high level of *iso*-OPDA upon *cis*-(±)-OPDA application (**Figure 6B**; **Supplemental Figure S5**). Moreover, we also observed that *iso*-OPDA is further metabolized in moss, likely conjugated to amino acids, since in the oxylipin profile (OPDA-Asp and OPDA-Glu) of the wild-type we observed peaks that were absent in the *gh3.2* single and *gh3* double knockout mutants (**Supplemental Figure S6**). The conifer *P. abies* possesses several GH3 proteins that fall within Group II, and we previously characterized four members conjugating auxins (Brunoni et al., 2020; 2023). We tested the ability of these recombinant PaGH3s to target *cis*-(±)-OPDA and (±)-JA, and only PaGH3.17 was observed to conjugate *cis*-(±)-OPDA primarily with Glu and to a lesser extent with Phe and Trp, while no conjugation of (±)-JA occurred with any of these recombinant PaGH3s (**Figure 6A**; **Supplemental Dataset S1**; **Supplemental Table S1**). Feeding spruce seedlings with *cis*-(±)-OPDA showed that it is mainly conjugated with Val and only to a lesser extent with Glu, Asp, Ile, Trp, and Phe. In contrast, no formation of OPDA-Ala was observed, demonstrating that conjugation of *cis*-OPDA occurs mainly with Val in spruce and suggesting that other than PaGH3.17 yet-to-be-characterized PaGH3s might be responsible for this reaction (**Figure 6C**). These results confirm that *cis*-OPDA amino acid conjugation is an evolutionarily conserved metabolic pathway in plants.

**Figure 6.**
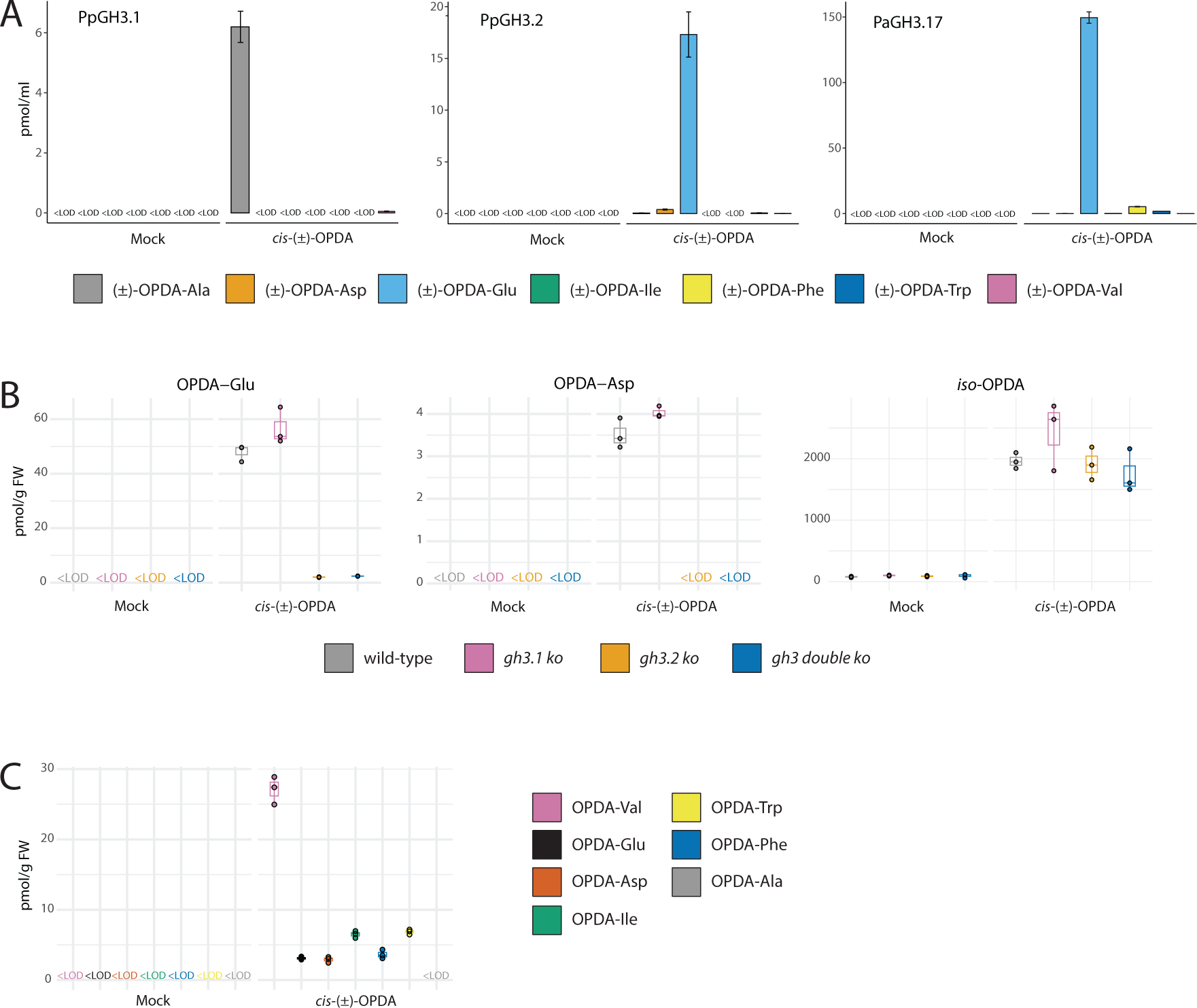
Amino acid conjugation of *cis*-OPDA is an evolutionarily conserved metabolic pathway in plants. A, Conjugating activity with *cis*-(±)-OPDA of recombinant *P. patens* (Pp) and *P. abies* (Pa) GH3s in the bacterial assay. Enzymatic assay and control reactions were carried out as reported in Figure 5A. (±)-OPDA-aa level is expressed as pmol/ml. B, Formation of OPDA-Glu, OPDA-Asp, and *iso*-OPDA after feeding of 3-week-old *P. patens* wild-type, *gh3.1*, *gh3.2* single and *gh3.1,gh3.2* double knockout mutant gametophores with or without 50 µM *cis*-(±)-OPDA for 24 h. C, Formation of indicated OPDA-aa after the feeding of 2-week-old *P. abies* seedlings with or without 50 µM *cis*-(±)-OPDA for 6 h. Metabolite concentrations are given as pmoles per gram fresh weight (FW). Horizontal lines in the box plots are medians, boxes show the upper and lower quartiles, and whiskers show the full data range. Mean ± SD (*n*=3). Below the limit of detection, <LOD.

## Discussion

Conjugation of *cis*-OPDA to amino acids has been reported to occur in angiosperms, as OPDA-Ile and OPDA-Asp were identified in Arabidopsis wounded leaves and in chitooligosaccharide-treated rice cell culture, respectively (Floková et al., 2016; Shinya et al., 2022). Although it was proposed that these conjugates might play a role in forming of *cis*-OPDA active ligands, the mechanism by which OPDA-aa are formed, their role and function in plants remain elusive and poorly understood. The popularization of mass spectrometry (MS) techniques combined with the use of isotope-labeled internal standards provides the means for accurate quantification and precise identification of plant metabolites independently of the original matrix (Brunoni et al., 2022). Here, we investigated the occurrence and meaning of *cis*-OPDA amino acid conjugation using a liquid chromatography (LC)-MS/MS-based method to study OPDA-aa formation under different physiological conditions that alter *cis*-OPDA and JA homeostasis in plants. Our results show that both biotic and abiotic stress elicits the conjugation of *cis*-OPDA to amino acids in Arabidopsis, very likely because of the *cis*-OPDA level increase (**Figure 1**; **Supplemental Figure 1**). We further demonstrated that these (±)-OPDA-aa possess *cis*-OPDA-like activity on typical transcriptional and physiological *cis*-OPDA- and JA-responses in Arabidopsis (**Figure 2**; **Figure 3A**) and that the (±)-OPDA-aa-mediated root-growth inhibition, similar to *cis*-(±)-OPDA and (±)-JA, requires functional conversion of JA into JA-Ile, thus possible conversion of OPDA-aa to *cis*-OPDA might be needed to exert their function (**Figure 2, A** and **B**). To address this latter question, we performed a feeding assay using isotopically labeled (±)-OPDA-Glu and (±)-OPDA-Val and confirmed that OPDA-Val is cleaved into *cis*-OPDA and Val by plants. In contrast, OPDA-Glu did not seem to follow the same fate, although we could not exclude a potential cleavage site different from the amide bond and/or Glu further metabolic conversion (**Figure 3B**). Further biochemical and physiological analyses were conducted to investigate the pathway for OPDA-aa formation and hydrolysis. We addressed the possible involvement of members of the GH3 and the ILR1/ILL families known to catalyze amino acid conjugation of acidic phytohormones and hydrolysis of the amino acid conjugates, respectively. GH3 proteins are canonically divided into three clades based on their hormone selectivity. Consistent with this classification, we investigated the possible participation of the JA-conjugating GH3 proteins in recognition of *cis*-OPDA, but surprisingly we could not confirm any involvement of Group I GH3s in an enzymatic assay or plants (**Supplemental Dataset S1**; **Supplemental Figure S2**). Further functional analysis of the other Arabidopsis GH3 enzymes combined with *in planta*-feeding assay of *gh3* sextuple mutant revealed that OPDA-Asp formation depends on the activity of the IAA-inactivating enzymes and that members of the Group III GH3s are likely involved in the conjugation of *cis*-OPDA with the other amino acids (**Figure 4**; **Supplemental Figure S3A**; **Supplemental Dataset S1**). Three enzymes, ILR1, ILL6, and IAR3, of the ILR1/ILL family were previously identified as JA-amidohydrolases that are induced upon wounding (Widemann et al., 2013; Zhang et al., 2016). Here, we investigated whether these JA-amidohydrolases and the additional member of the ILR1/ILL family, ILL2, mediated the cleavage of the OPDA-aa by performing a functional analysis coupled with *in vivo* study of their activity in wounding response. Recombinant amidohydrolases showed overlapping but distinct substrate specificities for various amino acid conjugates of *cis*-OPDA (**Figure 5A**; **Supplemental Dataset S1**). The wounding response of plants knocked-out for the four genes was consistent with ILR1, IAR3, ILL6, and ILL2 being the *cis*-OPDA amidohydrolases, thus demonstrating that in wounded leaves, these amidohydrolases contribute to regulating active *cis*-OPDA level (**Figure 5B**; **Supplemental Figure S4**). Considering the tissue-specific and developmentally controlled expression of *GH3* and *ILR1/ILL* genes (Rampey et al., 2004; http://bar.utoronto.ca/), OPDA-aa formation and hydrolysis may become more critical for localized *cis*-OPDA increase/decrease in specific cell types, thus, in addition to wounding or pathogen infection, *cis*-OPDA/OPDA-aa may play a role based on a regular developmental program. Furthermore, the levels of accumulated OPDA-aa in these responses are much less than that of JA-Ile, thus we have to consider homeostasis of OPDA-aa as an additional determinant in different stress responses and developmental processes.

Oxylipin biosynthesis in bryophytes and gymnosperms is significantly less studied than in flowering plants, thus impeding our understanding of these biosynthetic mechanisms in an evolutionary context. Therefore, this study further investigated the possible conservation of the *cis*-OPDA conjugative pathway in the moss *P. patens* and the conifer *P. abies*. Orthologs of critical enzymes for JA/JA-Ile biosynthesis appear absent in bryophytes but are present in gymnosperms (Chini et al., 2023). Consequently, bryophytes can synthesize the JA precursor *cis*-OPDA but they cannot produce JA/JA-Ile, whereas gymnosperms can synthesize JA/JA-Ile similarly to flowering plants. Consistently, we have recently shown that wounding induces *cis*-OPDA, JA, JA-Ile, and JA-Phe accumulation in spruce hypocotyl cuttings (Alallaq et al., 2020). Herein, results from enzymatic and feeding assays showed that *cis*-OPDA could be conjugated to various amino acids in spruce and that, similarly to Arabidopsis, the JA-conjugating GH3 enzymes differ from those accepting *cis*-(±)-OPDA (**Figure 6, A and C**; **Figure 4**; **Supplemental Figure S3**; **Supplemental Dataset S1**; **Supplemental Table S1**). Despite the lack of JA/JA-Ile, bryophytes and vascular plants share a conserved COI1 signaling machinery that is activated by distinct molecules, such as dnOPDA isomers or JA-Ile, respectively (Monte et al., 2020), thus demonstrating that *cis*-OPDA does not function as a COI1 ligand itself in bryophytes neither. Instead, *cis*-OPDA appears to be precursors of dnOPDA isomers in the liverwort *Marchantia* and *iso*-OPDA in the moss *P. patens* (Monte et al., 2018; Mukhtarova et al., 2020). Therefore, it seems unlikely that OPDA-aa could function as COI1 ligands, although a possible interaction between the COI1-JAZ co-receptor complex and any OPDA-aa was not investigated in this work. Finally, our findings from enzymatic and feeding assays demonstrated that GH3-mediated conjugation of *cis*-OPDA to amino acids is a conserved metabolic route in moss and confirmed that *cis*-OPDA is also a source of *iso*-OPDA that, in turn, might be further conjugated to amino acids itself (**Figure 6, A and B**; **Supplemental Figure, S5 and S6**). Although additional studies are needed to understand the biological function of synthesis and hydrolysis of OPDA-aa, this work suggests this pathway’s potential contribution and conservation in the temporary storage of *cis*-OPDA. Thus, *cis*-OPDA and other acidic phytohormones, such as IAA and JA, appear to share the same level of regulation that relies on similar metabolic machinery to modulate the homeostasis of the active phytohormone.

## Materials and methods

### Plant material

*Arabidopsis thaliana* ecotype Col-0 was used as wild-type for all the experiments, except for **Figure 5B and Supplemental Figure S4**, where Wassilewskija (Ws) was also used. Knockout lines in the Col-0 background used were *jar1-11* (SALK_034543), *gh3* sextuple (*gh3.1,gh3.2,gh3.3,gh3.4,gh3.5,gh3.6*; Porco et al., 2016), and *ill6-2* (SALK_024894) whereas in the Ws background were *ilr1-1,iar3-2,ill2-1* triple and corresponding single mutants (Bartel and Fink, 1995; Davies et al., 1999; Rampey et al., 2004). Gametophores from *Physcomitrium patens* (Hedw.) Mitt. wild-type and knockout lines (International Moss Stock Center IMSC#40207 GH3-1/B34, IMSC#40208 GH3-2/22, and IMSC#40209 GH3-doKO-A96; Ludwig-Müller et al., 2009) were obtained from IMSC (www.moss-stockcenter.org). *Picea abies* (L. Karst) seeds were provided by SkogForsk (Såvar, Sweden).

### Mechanical wounding

Arabidopsis plants were grown in soil under neutral-day conditions (12 h light/12 h dark) in cultivation chambers maintained at 21 °C, with a light intensity of approximately 150 μmol m^-1^ s^-1^ and 40-60% relative humidity. Wounding was conducted on fully expanded rosette leaves of 6-week-old plants by crushing across the midvein three times using forceps (Widemann et al., 2013). At increasing time points following mechanical damage, leaf samples were quickly harvested and flash-frozen in liquid nitrogen before storing at -80 °C until use. Wounded leaves were pooled from 3-4 plants for each sample, and the experiment was repeated three times.

### Fungal infection analysis

Arabidopsis Col-0 seeds were gas sterilized with chlorine gas (10 ml of HCl (35%) in 50 ml of bleach) for at least 2 h in an airtight box. Seeds were then sown under sterile conditions on Petri dishes containing ½ Murashige-Skoog (½MS) medium (2.15 g salts including vitamins per 1 l) with 1% sucrose and 0.5 g l^-1^ MES monohydrate at pH 5.7 and solidified with 0.8 % Gellan gum. Stratification was carried out for 3 d at 4 °C and then, plates were transferred to light at 22 ± 1 °C, under long-day conditions (16 h light/8 h dark; 100 µmol^-2^ s^-1^) and grown vertically. Wild-type *Botrytis cinerea* CCF2361 isolated from a garden strawberry (Department of Botany, Faculty of Science, Charles University, Czech Republic) was grown on potato-dextrose agar (PDA; HiMedia Laboratories) at 22 °C. After three weeks, spores were collected with sterile distilled water and filtered through glass wool (Sigma-Aldrich). Two-week-old *in vitro*-grown Arabidopsis seedlings were inoculated with a 10-µl drop of fungal spores (17.10*6 spores/ml); infected plants were harvested four days post-inoculation and flash-frozen in liquid nitrogen before storing at -80 °C until LC-MS/MS analysis. About sixteen infected Arabidopsis seedlings per replicate were used, and the experiment was repeated three times.

### Feeding experiments

Plant growth conditions and feeding experiments of Arabidopsis, *P. abies* seedlings, and *P. patens* gametophores were carried out as described in Brunoni et al. (2023). For feeding with unlabeled standards, seven-day-old Arabidopsis seedlings were cultivated in liquid ½ MS medium supplemented with or without 50 µM (±)-JA, *cis*-(±)-OPDA, (±)-OPDA-Ala, (±)-OPDA-Val, (±)-OPDA-Phe, (±)-OPDA-Asp, (±)-OPDA-Glu, and (±)-OPDA-Phe for 3 h. Spruce seedlings were cultivated in liquid ½ MS medium supplemented with or without 50 µM *cis*-(±)-OPDA for 6 h. Moss gametophores were cultivated in liquid Knopp medium supplemented with or without 50 µM *cis*-(±)-OPDA for 24 h.

For feeding with labeled standards, seven-day-old Arabidopsis seedlings were cultivated in liquid ½ MS medium supplemented with 50 µM *cis*-(±)-OPDA-[^13^C ^15^N]*L*-glutamate and *cis*-(±)-OPDA-[^13^C ^15^N]*L*-valine. One ml of liquid medium and 10 mg FW of plant samples were collected after 30 min, 1, 3, 6, and 24 h. The feeding experiments were repeated three times.

### Root-growth inhibition assay

Arabidopsis Col-0 and *jar1-11* seeds were germinated and vertically grown in the conditions described above. Root length of 20-25 seedlings was measured seven days after germination, alone or in the presence of 50 µM (±)-JA, 5 µM *cis*-(±)-OPDA, (±)-OPDA-Ala, (±)-OPDA-Val, (±)-OPDA-Phe, (±)-OPDA-Asp, (±)-OPDA-Glu, and (±)-OPDA-Phe. Plates were scanned using a densitometer (GS-800, Bio-Rad), and root length measurement was performed with the Simple Neuronal Tracing plugin of ImageJ/Fiji software. Three independent biological replicates were measured for each sample. Data were analyzed by one-way ANOVA/Tukey HSD *post hoc* test (*P*<0.05).

### Promoter-GUS assay

Arabidopsis seeds of *35S:JAZ1-GUS* and *35S:JAZ9-GUS* marker lines (Thines et al., 2007) were germinated and vertically grown in the conditions described above. Seven-day-old *in vitro*-grown seedlings were treated with or without 10 µM (±)-JA, *cis*-(±)-OPDA, (±)-OPDA-Ala, (±)-OPDA-Val, (±)-OPDA-Phe, (±)-OPDA-Asp, (±)-OPDA-Glu, and (±)-OPDA-Phe for 2 h and the visualization of GUS was carried out as described in Chini et al. (2018). Root imaging was carried out with a camera (QuickPHOTO CAMERA 3.2) connected to a stereo microscope (Olympus DP72).

### RNA extraction and real-time qPCR

Total RNA was isolated using Spectrum Total RNA kit (Sigma-Aldrich), and DNA-free™ DNA Removal Kit (Invitrogen) was used to prepare DNA-free RNA according to the manufacturer’s instructions. One µg of total RNA was reverse transcribed for each sample with RevertAid H Minus Reverse Transcriptase (Thermo Scientific). Quantitative real-time PCR (RT-qPCR) was performed on a CFX384 Touch Real-Time PCR Detection System (Bio-Rad) using 2× SYBR Green Real-Time PCR Master Mix (Applied Biosystems). The three-step cycling program was as follows: 95 °C for 2 min, followed by 40 cycles at 95 °C for 5 s, 60.5 °C for 20 s and 72 °C for 10 s. Melting curve analysis was conducted between 75 and 95 °C. The *ACT2* (AT3G18780) gene was used as a constitutive internal standard to normalize the obtained gene expression results. Primer sequences are listed in **Supplemental Table S2**. Expression levels were calculated using the ΔΔCt method (Pfaffl, 2001). Three biological replicates were performed for each test. Data were analyzed by one-way ANOVA/Tukey HSD *post hoc* test (*P*<0.05).

### Cloning, protein production, and bacterial enzyme assay

*Escherichia coli* BL21 (DE3) strains expressing recombinant AtGH3s, PpGH3s, and PaGH3s used in this work were previously generated (Brunoni et al., 2019b; 2020; 2023). Recombinant protein production and enzymatic assay of AtGH3s, PpGH3s, and PaGH3s were performed as described previously (Brunoni et al., 2023). AtIAR3, AtILL2, AtILL6, and AtILR1 open reading frame sequences deleted of the 25-N-terminal signal peptide-encoding codons were amplified using *Phusion* Taq Polymerase (Thermo Fisher) prior to cloning into pETM11 plasmid in the BL21 (DE3) *E. coli* strain. Recombinant protein production was performed as described by Brunoni et al. (2019b). For conjugation assay, 500 µl of clarified cell lysate from GH3-producing bacterial cultures was incubated with GH3 cofactors and with or without 0.1 mM (±)-JA and *cis*-(±)-OPDA for 5 h at 30 °C with constant shaking at 50 rpm in darkness. For hydrolysis assay, 500 µl of clarified cell lysate from amidohydrolase-producing bacterial cultures was incubated with 1 mM MgCl_2_ and 0.1 mM IAA-Ala, (±)-OPDA-Ala, (±)-OPDA-Val, (±)-OPDA-Phe, (±)-OPDA-Asp, (±)-OPDA-Glu, and (±)-OPDA-Ile in the same conditions as those for the conjugation assay.

### Chemical synthesis and liquid chromatography tandem mass spectrometry phytohormone measurements

Detailed procedures for synthesis and phytohormone measurements are reported in **Supplemental Note 1**.

## Acknowledgements

Plant Sciences Core Facility of CEITEC Masaryk University is acknowledged for the technical support. We thank Hana Svobodová, Veronika Večeřová, and Miroslava Špičáková for technical support. We also thank Andrea Chini (Universidad Autónoma de Madrid, Spain) for providing *JAZ-GUS* lines, Thierry Heitz (Institut de Biologie Moléculaire des Plantes du CNRS, France) for *ill6-2* mutant seeds, Bonnie Bartel (Rice University, USA) for *ilr1-1*, *iar3-2*, *ill2-1* single and triple mutant seeds, and Paul Staswick (University of Nebraska–Lincoln, USA) for *jar1-11* mutant seeds.

We apologize to those whose work could not be cited because of space constraints.

## Author contributions

FB and ON conceived the study. FB designed the experiments. FB, JŠ, VM, TP, MKr, AA, MP and MKa performed the experiments. MU synthesized the *iso*-OPDA standard. MH performed the statistics and data visualization. KF and MU acquired the financial support. CW and MS contributed to the discussion. FB wrote the paper with input from all the authors.

## Funding

This work was financially supported by Czech Science Foundation project No. 19-10464Y (to KF), and by Grant-in-Aid for Scientific Research from JSPS (Japan) projects No. 23H00316, 23H04883, and JPJSBP120239903 (to MU).

## Conflict of interest

On behalf of all authors, the corresponding author states that there is no conflict of interest.

## Supplemental Data

**Supplemental Figure S1** Accumulation of *cis*-OPDA, JA, and JA-aa in Arabidopsis plants during stress responses. A, Time-course accumulation of *cis*-OPDA, JA, and indicated JA-aa in Col-0 after leaf wounding. Six-week-old plants were wounded, and damaged leaves were collected after the indicated times. B, Accumulation of *cis*-OPDA, JA, and indicated JA-aa in Col-0 plants infected with *B. cinerea* 3 days after inoculation. Metabolite levels are expressed as pmoles per gram fresh weight (FW). Mean ± SD (*n*=3). Below the limit of detection, <LOD.

**Supplemental Figure S2** Amino acid conjugation of *cis*-OPDA is not mediated by JAR1/GH3.11 *in planta*. Accumulation of indicated JA-aa and OPDA-aa after exogenous treatment with or without 50 µM (±)-JA and *cis*-(±)-OPDA in *jar1-11* mutants. Asterisk indicates statistically significant differences, as determined by Student’s *t*-test (*P*<0.05). Metabolite concentrations are given as pmoles per gram dry weight (DW). Mean ± SD (*n*=3). Below the limit of detection, <LOD.

**Supplemental Figure S3** Conjugating activity with *cis*-(±)-OPDA of recombinant AtGH3s and accumulation of OPDA-aa in *gh3* sextuple mutant upon *cis*-(±)-OPDA feeding. A, Analysis of OPDA-aa synthesized by recombinant AtGH3.12, AtGH3.14, AtGH3.15, and AtGH3.17 in the bacterial assay. The cell lysate was incubated with or without 0.1 mM *cis*-(±)-OPDA and GH3 cofactor mixture for 5 h at 30 °C. The bacterial assay carried out with cell lysate from GFP-producing bacteria was used as a negative control. Cell lysate without *cis*-(±)-OPDA and cofactor mixture was used as a mock sample. OPDA-aa level is expressed as pmol/ml. B, Formation of OPDA-Glu, OPDA-Ala, OPDA-Val, OPDA-Ile, and OPDA-Phe after feeding of 7-day-old Arabidopsis Col-0 and *gh3* sextuple mutant (*gh3.1,gh3.2,gh3.3,gh3.4,gh3.5,gh3.6*) with or without 50 µM *cis*-(±)-OPDA for 3 h. OPDA-aa concentration is given as pmoles per gram fresh weight (FW). Horizontal lines in the box plots are medians, boxes show the upper and lower quartiles, and whiskers show the entire data range. Mean ± SD (*n*=3). Below the limit of detection, <LOD.

**Supplemental Figure S4** Accumulation of OPDA-aa in *ilr/ill* single knockout mutants upon wounding. Time-course accumulation of indicated OPDA-aa in Ws, *ilr-1*, *iar3-2*, and *ill2-1* single knockout mutants after leaf wounding. Six-week-old plants were wounded, and damaged leaves were collected after the indicated times. Asterisk indicates statistically significant differences, as determined by Student’s *t*-test (Ws vs *ilr1-1, iar3-2,* or *ill2-1*; *P*<0.05). OPDA-aa concentrations are given as pmoles per gram fresh weight (FW). Mean ± SD (*n*=3). Below the limit of detection, <LOD.

**Supplemental Figure S5** Extracted ion chromatogram of *cis*-/*iso*-OPDA and their identification in *Physcomitrium patens* after feeding with *cis*-(±)-OPDA (50 µM, 24 h) wild-type gametophores.

**Supplemental Figure S6** Extracted ion chromatogram of OPDA-Asp and OPDA-Glu standard (A) and *Physcomitrium patens* wild-type (B), *gh3.1* single (C), *gh3.2* single (D) and *gh3* double (E) knockout mutant gametophores after feeding with *cis*-(±)-OPDA (50 µM, 24 h).

**Supplemental Table S1** Semi-quantitative comparison of enzymatic activities with (±)-JA and amino acids of recombinant *A. thaliana*, *P. abies* and *P. patens* GH3 proteins resulting from the bacterial assay.

**Supplemental Table S2** List of primers sequences used for qPCR analysis and cloning.

**Supplemental Dataset S1** Raw mass spectrometry data of (±)-OPDA-aa and (±)-JA-aa with recombinant GH3 enzymes, and *cis*-(±)-OPDA and IAA with recombinant ILR1/ILL enzymes from the bacterial assay.

**Supplemental Note S1** Synthetic procedures and phytohormone measurements.

## Notes

### Competing Interest Statement

The authors have declared no competing interest.

