## Supplemental Note S1 for "Conjugation of *cis*-OPDA with amino acids is a conserved pathway affecting *cis*-OPDA homeostasis upon stress responses"

*- Synthesis of labeled compounds cis-(±)-OPDA-[$^{13}C_5$,$^{15}N$]L-glutamate and cis-(±)-OPDA-[$^{13}C_5$,$^{15}N$]L-valine*

*Chemicals*. *cis*-(±)-OPDA was prepared according to Zimmerman and Feng (1978) and Mik et al. (2023), and labeled amino acids [$^{13}C_5$,$^{15}N$]*L*-glutamic acid and [$^{13}C_5$,$^{15}N$]*L*-valine were purchased from Cambridge Isotope Laboratories (Tewksbury, MA, USA). A solution of isobutyl carbonic anhydride of *cis*-(±)-OPDA (*cis*-(±)-OPDA 1 eq., IBCF 1.1 eq., TEA 1.2 eq., THF, -10 °C, 30 min, filtration using MicroSpin centrifugation filter NYLON 0.2 µm) was added under nitrogen atmosphere to the solution of a silylated amino acid (amino acid 3 eq., TMS-Cl 7.5 eq. for Val and 10.5 eq. for Glu, TEA 12 eq., MeCN, reflux until dissolution) cooled to 0 °C. The resulting mixture was stirred at 0 °C for 2 h. After acidification by 1M HCl to pH ~ 3, the mixture was diluted with water and extracted by $CHCl_3$ (4× 5 ml). Combined organic layers were washed with brine, dried over $Na_2SO_4$, filtered, and evaporated under reduced pressure. Products were purified using preparative HPLC. Fractions containing purified products were partially evaporated and extracted by $CHCl_3$. Combined organic layers were dried over $Na_2SO_4$, filtered, and evaporated under reduced pressure.

*Analytical methods*. Purification of compounds – 1290 Infinity II LC/MSD preparative system (Agilent, Santa Clara, CA, USA) using 5 Prep-C18 column (100 × 21.2 mm, particle size 5 µm, Agilent) with a flow rate 20 ml/min using following gradients *cis*-(±)-OPDA-[$^{13}C_5$,$^{15}N$]*L*-Val: 0.1% AcOH (A), MeCN (B), gradient – 0 min 50% A, 3.5 min 50% A, 7 min 10% A, 7.5 min 10% A; *cis*-(±)-OPDA-[$^{13}C_5$,$^{15}N$]*L*-Glu: 0.1% AcOH (A), MeOH (B), gradient – 0 min 50% A, 8 min 10% A, 8.5 min 10% A. Compound purity and mass spectra - AQUITY UPLC® H-Glass system (Waters, Milford, MA, USA) using Symmetry C18 column (150 × 2.1 mm, particle size 5 µm, Waters) linked to ACQUITY UPLC PDA detector and single quadrupole mass spectrometer QDa (Waters). NMR spectra – JEOL ECZ/R-400 spectrometer (Jeol, Japan), HRMS analysis – SYNAPT G2-Si (Waters, Milford, MA, USA).

*Analytical data for labelled compounds.*

**{8-[(1S*,5S*)-4-oxo-5-((Z)-pent-2-en-1-yl)cyclopent-2-en-1-yl]octanoyl}-*L*-glutamic-[$^{13}C_5$,$^{15}N$] acid**

Colorless gel, yield: 56%. HPLC-UV/VIS retention time, purity (min., %): 22.20, >98%. ESI$^+$-MS *m/z* (rel. int. %, ion): 428.5. $^1H\{^{13}C\}$-NMR (400 MHz, $CDCl_3$) δ (ppm): 0.95 (t, *J* = 7.5 Hz, 3H), 1.14-1.20 (m, 1H), 1.29 (s, 8H), 1.60-1.71 (m, 3H), 2.00-2.16 (m, 4H), 2.26 (t, *J* = 7.1 Hz, 3H), 2.43-2.50 (m, 4H), 2.98 (s, 1H), 4.60 (s, 1H), 5.34-5.43 (m, 2H), 6.19 (d, *J* = 5.7 Hz, 1H), 6.77 (d, *J* = 6.6 Hz, 1H), 7.00 (d, *J* = 6.4 Hz, 1H), 7.10 (s, 2H), 7.77 (dd, J = 5.5, 2.3 Hz, 1H). $^{13}C$-NMR (100 MHz, $CDCl_3$) δ (ppm): 14.0, 20.7, 23.7, 25.4, 26.8 (t, *J* = 34.6 Hz), 27.4, 29.0, 29.1, 29.9 (dd, *J* = 53.9, 34.0 Hz), 30.6, 44.3, 49.9, 51.2-52.0 (m), 126.8, 132.2, 133.1, 168.1, 174.8 (d, *J* = 57.8 Hz), 177.16 (d, *J* = 55.8 Hz), 211.9. HRMS (ESI-TOF): *m/z* calc for $C_{18}{}^{13}C_5H_{35}{}^{15}NO_6$ $[M+H]^+$ 428.2675, found 428.2685.

**{8-[(1S*,5S*)-4-oxo-5-((Z)-pent-2-en-1-yl)cyclopent-2-en-1-yl]octanoyl}-*L*-valine-[$^{13}C_5$,$^{15}N$] acid**

Colorless gel, yield: 34%. HPLC-UV/VIS retention time, purity (min., %): 24.54, >98%. ESI$^+$-MS *m/z* (rel. int. %, ion): 398.5. $^1H\{^{13}C\}$-NMR (400 MHz, $CDCl_3$) δ (ppm): 0.89-0.98 (m, 9H), 1.14-1.19 (m, 1H), 1.30-1.40 (m, 8H), 1.63-1.75 (m, 3H), 2.03-2.17 (m, 3H), 2.24-2.33 (m, 3H), 2.46-2.52 (m, 2H), 2.99 (bs, 1H), 4.58 (bs, 1H), 5.37-5.41 (m, 2H), 5.94-5.96 (m, 1H), 6.19 (bs, 1H), 7.75 (bs, 1H). $^{13}C$-NMR (100 MHz, $CDCl_3$) δ (ppm): 14.0, 17.6 (d, *J* = 34.7 Hz), 19.0 (d, *J* = 34.7 Hz), 20.8, 23.7, 25.6, 27.5, 27.5, 29.0, 29.1, 29.6, 30.8 (q, *J* = 34.8 Hz), 36.5, 36.6, 44.3, 49.9, 56.4-57.3 (m), 126.9, 132.4, 133.0, 167.5, 167.5, 174.9 (d, *J* = 57.8 Hz), 211.3, 211.4. HRMS (ESI-TOF): *m/z* calc for $C_{18}{}^{13}C_5H_{37}{}^{15}NO_4$ $[M+H]^+$ 398.2933, found 398.2937.

*- Phytohormone measurements*

*Chemicals*. Authentic and stable isotope labeled analytical standards were purchased from OlChemIm Ltd. (Olomouc, Czech Republic): *cis*-(±)-OPDA, (±)-JA, (-)-JA-Val, (-)-JA-Ile, (-)-JA-Phe, [$^2H_5$]OPDA, [$^2H_6$]JA, [$^2H_2$]JA-Ile, [$^{13}C_6$]IAA; IAA was purchased from Sigma Aldrich (St. Louis, MO, USA); [$^2H_5$]OPDA-Ile was synthesized as described by Zimmerman and Feng (1978) and Krammel et al. (1988); (±)-JA-Ala, (±)-JA-Asp, (±)-JA-Gly, (±)-JA-Glu, (-)-JA-Met and (±)-JA-Trp were synthetised as described by Kramell et al. (1988) and *iso*-OPDA was synthesized as described by Inagaki et al. (2021). All the chemicals used for sample preparation and analysis were highly analytical.

The samples from bacterial enzyme assays (1 ml) were centrifuged (20 000 *g*, 10 min, 8 °C), 2 µl of the supernatant and 10 µl of IS mixture (10 pmol of [$^2H_6$]JA, 5 pmol of [$^2H_2$]JA-Ile, [$^2H_5$]OPDA, [$^2H_5$]OPDA-Ile and 5 pmols [$^{13}C_6$]IAA) were added into 28 µl of 20% aqueous acetonitrile. The samples were mixed with a pipette tip and analyzed by a liquid chromatography-tandem mass spectrometry system (LC-MS/MS).

The frozen plant material was powdered under liquid nitrogen with mortar and pestle and weighed in samples of 10 mg FW into 2 ml Eppendorf tubes. To each sample, 1 ml of 50% aqueous methanol, a mixture of IS (10 pmols of [$^2H_6$]JA, 5 pmols [$^2H_2$]JA-Ile, [$^2H_5$]OPDA and [$^2H_5$]OPDA-Ile) and 4 ceria-stabilized zirconium oxide 2 mm beads (Retsch GmbH, Haan, Germany) were added. The samples were homogenized on a MM 400 mixer mill (Retsch GmbH, Haan, Germany) (27 Hz, 6 min, precooled holders) and centrifuged (25 800 *g*, 15 min, 4 °C). The supernatants were purified on solid phase extraction columns Oasis® HLB 1cc 30 mg columns (Waters, Milford, MA, USA) as described by Mik et al. (2023).

All samples were analyzed on an Agilent 6490 Triple Quadrupole LC/MS system coupled to a 1290 Infinity LC system (Agilent Technologies, Santa Clara, CA, USA) using chromatographic conditions and MS/MS parameters described by Široká et al. (2022) and Mik et al. (2023) and reported in **Supplemental Note Table 1** and **Supplemental Note Figure 1**. The elution order of (+) and (-) JA-aa diastereomers was specified following Kramell et al. (1988) and Jikumaru et al. (2004). (±)-JA-Gly, (±)-JA-Glu, (±)-JA-Asp, (±)-JA-Trp were quantified as sum (Σ) of unresolved or partly resolved peaks. (±)-JA-Asp and (±)-JA-Glu presence in the samples was identified based on the agreement of their RT and rates of confirmatory and reference MRMs with corresponding analytical standards (**Supplemental Note Figure 2**). Also, for other JA-aa and OPDA-aa identification, these criteria were applied. The *cis*- and *iso*-OPDA were identified in *Physcomitrium patens* based on their RT corresponding to the analytical standard (**Supplemental Figure S5**). The separation of the compounds analyzed is depicted in the **Supplemental Note Figure 3.**
The MS system was operated in dynamic multiple reaction monitoring mode in positive and negative electrospray ionization mode. The nozzle voltage was set to 0 V, the capillary voltage to 2800/3000 V positive/negative mode, and the drying gas was at 130 °C with a flow rate of 14 l/min. The sheath gas was heated to 400 °C, and its flow rate was 12 l/min. The MassHunter Quantitative software package version B.09.00 (Agilent Technologies, Santa Clara, CA, USA) was used for data processing. The levels of the analytes were determined using a calibration curve prepared in the range of 0.0045 – 4.5 pmol for compounds in positive ionization and 0.0045 – 45 pmol for JA (negative ionization), logarithmically scaled.

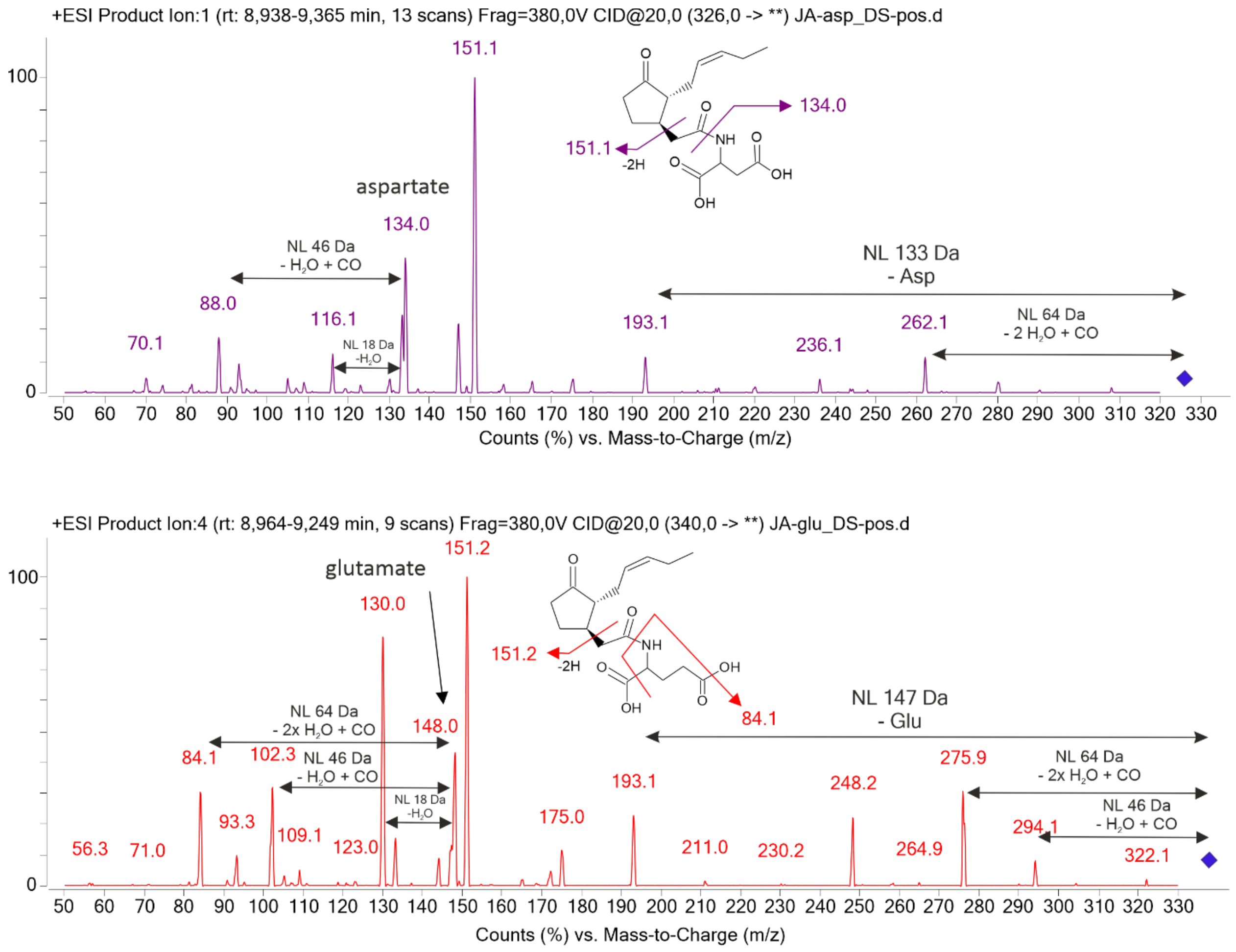


**Supplemental Note Figure 1** Fragmentation spectra of (±)-JA-Asp and (±)-JA-Glu at CE 20 eV.

**Supplemental Note Table 1** LC-MS/MS method parameters.

| Analyte | IS | Ionisation | Reference MRM (CE; eV) | Confirmatory MRM (CE; eV) | RT; min |
|---|---|---|---|---|---|
| *cis*-(±)-OPDA | [$^{2}H_5$]OPDA | $[M+H]^+$ | 293.2 > 275.2 (12) | | 13.7 |
| *iso*-OPDA | [$^{2}H_5$]OPDA | $[M+H]^+$ | 293.2 > 275.2 (12) | | 13.5 |
| IAA | [$^{13}C_6$]IAA | $[M+H]^+$ | 176.1 > 130.1 (24) | | 6.9 |
| (±)-JA | [$^{2}H_6$]JA | $[M-H]^-$ | 209.2 > 58.8 (8) | | 10.1 |
| (-)-JA-Ala/(+)-JA-Ala | [$^{2}H_2$]JA-Ile | $[M+H]^+$ | 282.1 > 151.1 (12) | 282.1 > 90.1 (16) | 9.5/9.7 |
| (±)-JA-Asp | [$^{2}H_2$]JA-Ile | $[M+H]^+$ | 326.1 > 151.1 (14) | 326.1 > 134.0 (16) | 9.2 |
| (±)-JA-Gly | [$^{2}H_2$]JA-Ile | $[M+H]^+$ | 268.1. > 151.1 (12) | 268.1 > 76.2 (16) | 8.8 |
| (±)-JA-Glu | [$^{2}H_2$]JA-Ile | $[M+H]^+$ | 340.2 > 151.1 (20) | 340.2 > 84.1 (40) | 9.1 |
| (-)-JA-Ile | [$^{2}H_2$]JA-Ile | $[M+H]^+$ | 324.3 > 151.2 (16) | 324.3 > 86.0 (26) | 12.2 |
| (-)-JA-Met | [$^{2}H_2$]JA-Ile | $[M+H]^+$ | 342.2 > 151.2 (20) | 342.2 > 193.0 (10) | 11.4 |
| (-)-JA-Phe | [$^{2}H_2$]JA-Ile | $[M+H]^+$ | 358.8 > 151.2 (16) | 358.8 > 120.1 (30) | 12.6 |
| (±)-JA-Trp | [$^{2}H_2$]JA-Ile | $[M+H]^+$ | 397.3 > 351.3 (12) | 397.2 > 151.0 (18) | 12.1 |
| (-)-JA-Val | [$^{2}H_2$]JA-Ile | $[M+H]^+$ | 310.3 > 151.3 (16) | 310.2 > 72.1 (30) | 11.4 |
| (±)-OPDA-Ala | [$^{2}H_5$]OPDA-Ile | $[M+H]^+$ | 369.3 > 280.0 (22) | 364.3 > 90.0 (20) | 13.4 |
| (±)-OPDA-Asp | [$^{2}H_5$]OPDA-Ile | $[M+H]^+$ | 408.3 > 275.3 (22) | 408.3 > 134.1 (22) | 13.3 |

| | | | | | |
|---|---|---|---|---|---|
| (±)-OPDA-Glu | [$^2H_5$]OPDA-Ile | $[M+H]^+$ | 422.3 > 275.3 (22) | 422.3 > 148.1 (22) | 13.0 |
| (±)-OPDA-Ile | [$^2H_5$]OPDA-Ile | $[M+H]^+$ | 406.2 > 86.1 (36) | 406.2 > 275.1 (28) | 14.4 |
| (±)-OPDA-Phe | [$^2H_5$]OPDA-Ile | $[M+H]^+$ | 440.3 > 120.1 (40) | 440.3 > 275.2 (30) | 14.8 |
| (±)-OPDA-Trp | [$^2H_5$]OPDA-Ile | $[M+H]^+$ | 479.2 > 158.9 (28) | 479.2 > 433.2 (22) | 14.3 |
| (±)-OPDA-Val | [$^2H_5$]OPDA-Ile | $[M+H]^+$ | 392.3 > 72.2 (40) | 392.3 > 275.1 (24) | 14.1 |

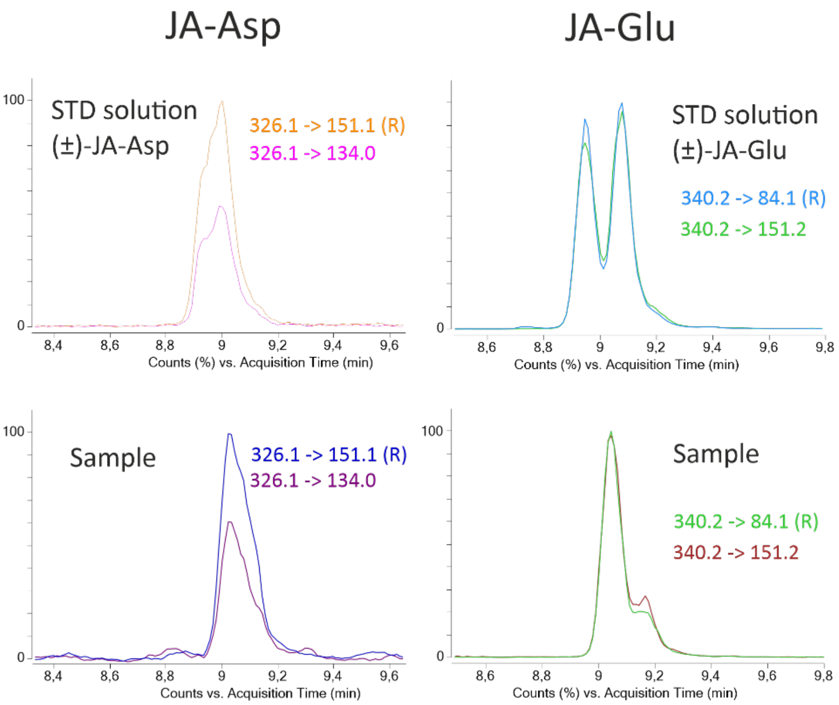


**Supplemental Note Figure 2** The extracted ion chromatogram of (±)-JA-Asp and (±)-JA-Glu and their identification in a sample.

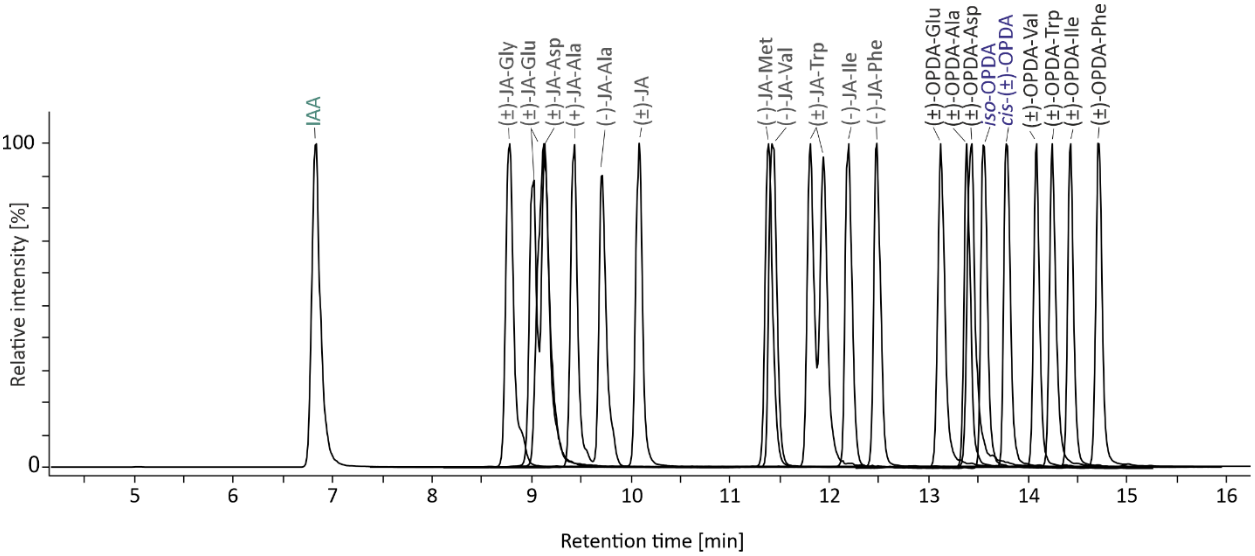


**Supplemental Note Figure 3** Chromatographic separation of IAA, (±)-JA, *cis*-(±)-OPDA, *iso*-OPDA, (±)-JA-aa and (±)-OPDA-aa.

- *Quantification of [$^{13}C_5$,$^{15}N$]L-glutamate and [$^{13}C_5$,$^{15}N$]L-valine*

The frozen plant material was powdered under liquid nitrogen with mortar and pestle and weighed into 10 mg of FW samples in 2 ml Eppendorf tubes. Each sample was extracted in 0.5 ml of 30% aqueous acetonitrile with the addition of internal standard - IS (10 pmol of [$^{13}C_4$,$^{15}N$]*L*-aspartate) and 3 ceria-stabilized zirconium oxide 2 mm beads (Retsch GmbH, Haan, Germany). The samples were homogenized on a MM 400 mixer mill (Retsch GmbH, Haan, Germany) (27 Hz, 6 min, precooled holders) and then centrifuged (25 800 *g*, 15 min, 4 °C). The 0.25 ml of supernatant was evaporated to dryness *in vacuo* Centrivap concentrator (Labconco, Kansas City, MO, USA) and derivatized using 14 µl of AccQ-Tag™ Ultra 2A "3X" Reagent and 4 µl of AccQ-Tag™ Ultra 1 Borate buffer (Waters, Milford, MA, USA). The analysis was performed on an LC-MS/MS Agilent 6490 Triple Quadrupole LC/MS with an electrospray coupled to a 1290 Infinity LC system (Agilent Technologies, Santa Clara, CA, USA), injecting 1 µl of each sample. The chromatography was performed at 53 °C on a SecurityGuard column-protected Kinetex Polar C18 reverse phase column 2.1 × 150 mm, particle size 2.6 µm (Phenomenex, Torrance, CA, USA). The mobile phases comprised 0.03% formic acid in water and acetonitrile at a flow rate of 0.53 ml/min. The chromatographic gradient started with 8 min long gradient from 1 to 8.5% acetonitrile, followed by 1 min long gradient from 8.5 to 10% of acetonitrile, continuing with 6 min long gradient from 10 to 12% of acetonitrile and 5 min long gradient from 12 to 98 % of acetonitrile. The acetonitrile content dropped from 98% to 1% within 1 min and was maintained for another 2 min before a new injection. The chromatographic separation of the analytes and the IS is shown in **Supplemental Note Figure 4**.
The LC-MS/MS parameters (*e.g.*, multiple reaction monitoring transitions (MRM), collision energies (CE)) were optimized using derivatized ([$^{13}C_4$,$^{15}N$]*L*-aspartate, [$^{13}C_5$,$^{15}N$]*L*-glutamate, [$^{13}C_5$,$^{15}N$]*L*-valine) and not derivatized (*cis*-(±)-OPDA-[$^{13}C_5$,$^{15}N$]*L*-glutamate, *cis*-(±)-OPDA-[$^{13}C_5$,$^{15}N$]*L*-valine) standards (**Supplemental Note Figures 5-8, Supplemental Note Table 2**). The MS system was operated in dynamic multiple reaction monitoring mode in positive electrospray ionization mode. The nozzle voltage was set to 0 V, the capillary voltage to 2800 V in positive mode, and the drying gas was at 160 °C with a flow rate of 14 l/min. The sheath gas was heated to 390 °C, and its flow rate was 12 l/min. The MassHunter Quantitative software package version B.09.00 (Agilent Technologies, Santa Clara, CA, USA) was used for data processing. The levels of the analytes were determined using a calibration curve prepared in the range of 0.001 – 10 pmol, logarithmically scaled.

**Supplemental Note Table 2** LC-MS/MS method parameters.

| Analyte | Diagnostic MRM | CE; eV | RT[a]; min |
|---|---|---|---|
| [$^{13}C_4$,$^{15}N$]*L*-aspartate – derivatized (IS) | 309.0-171.0 | 18 | 5.13 ± 0.02 |
| [$^{13}C_5$,$^{15}N$]*L*-glutamate - derivatized | 324.2-170.9 | 34 | 5.91 ± 0.02 |
| [$^{13}C_5$,$^{15}N$]*L*-valine – derivatized - | 294.2-171.0 | 26 | 12.06 ± 0.03 |
| *cis*-(±)-OPDA-[$^{13}C_5$,$^{15}N$]*L*-glutamate | 428.1-275.2 | 12 | 18.89 ± 0.01 |
| *cis*-(±)-OPDA-[$^{13}C_5$,$^{15}N$]*L*-valine | 398.2-77.0 | 18 | 19.53 ± 0.01 |

[a] RT – retention time, means ± standard deviation, *n* = 10.

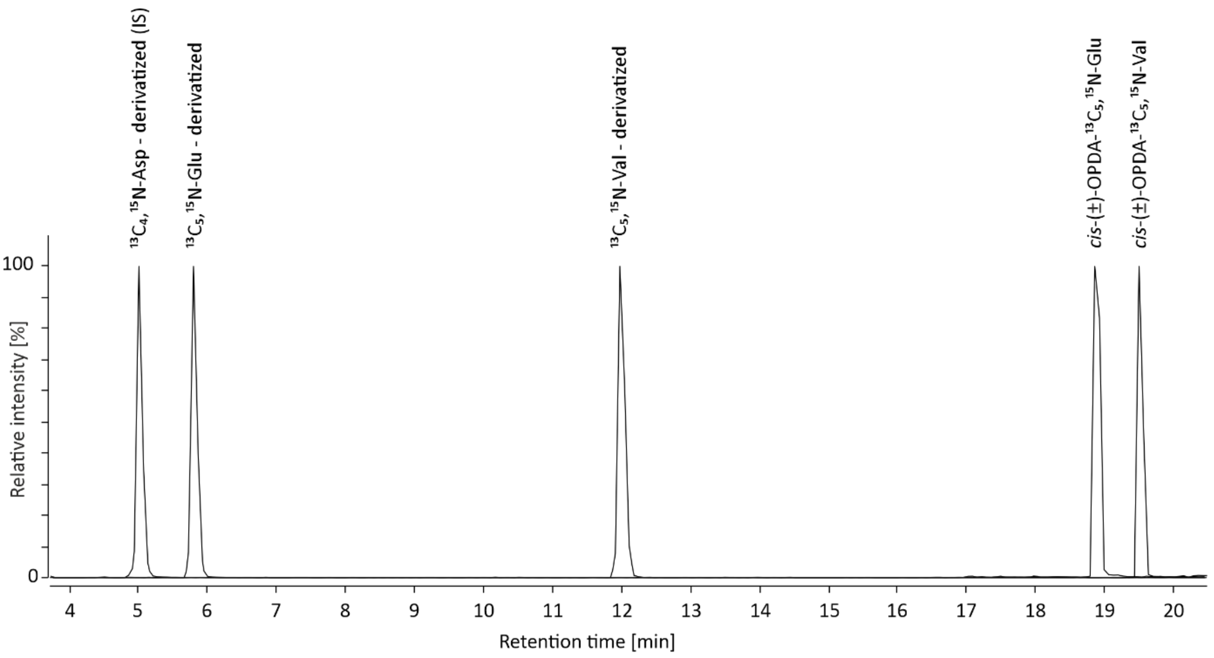


**Supplemental Note Figure 4** Chromatographic separation of analytes and IS.

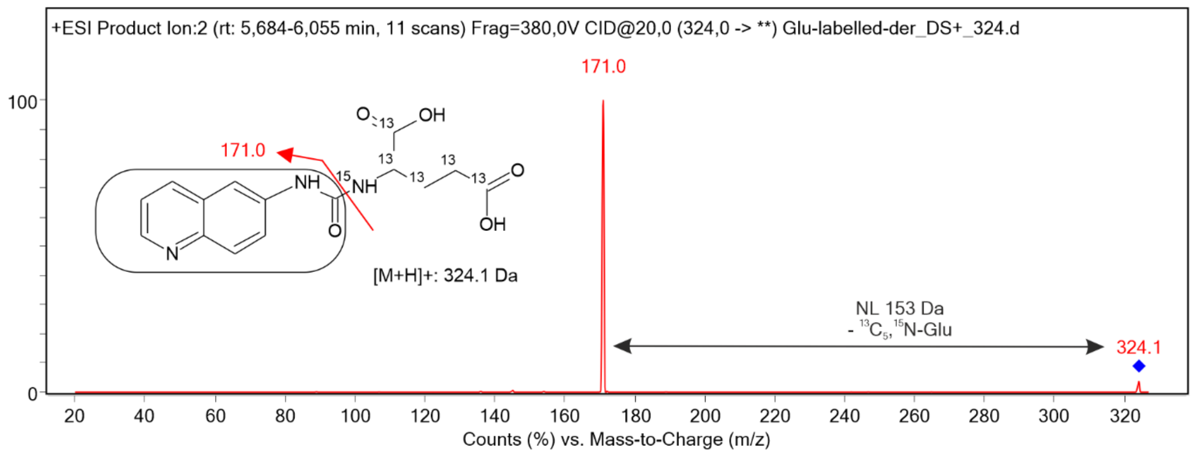


**Supplemental Note Figure 5** Fragmentation spectra at CE 20 eV of derivatized [$^{13}C_5$,$^{15}$N]*L*-glutamate.

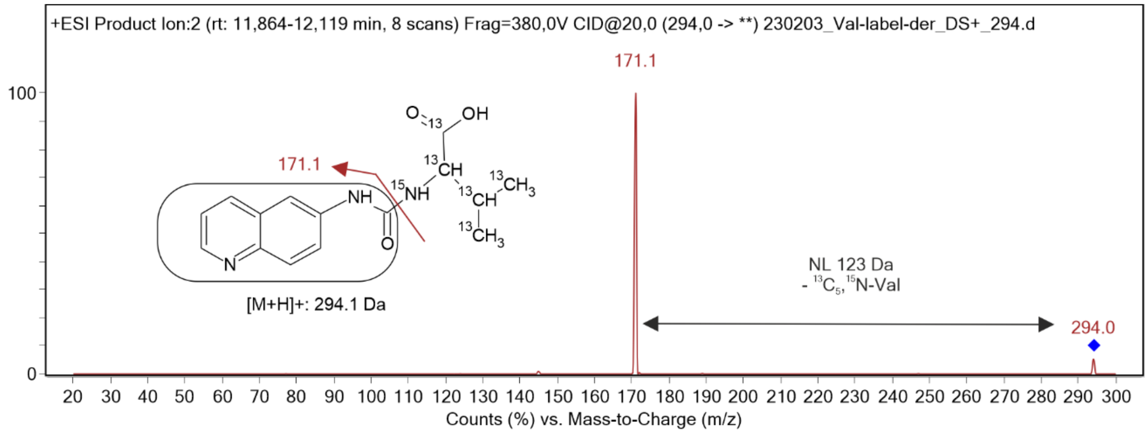


**Supplemental Note Figure 6** Fragmentation spectra at CE 20 eV of derivatized [$^{13}C_5$,$^{15}N$]*L*-valine.

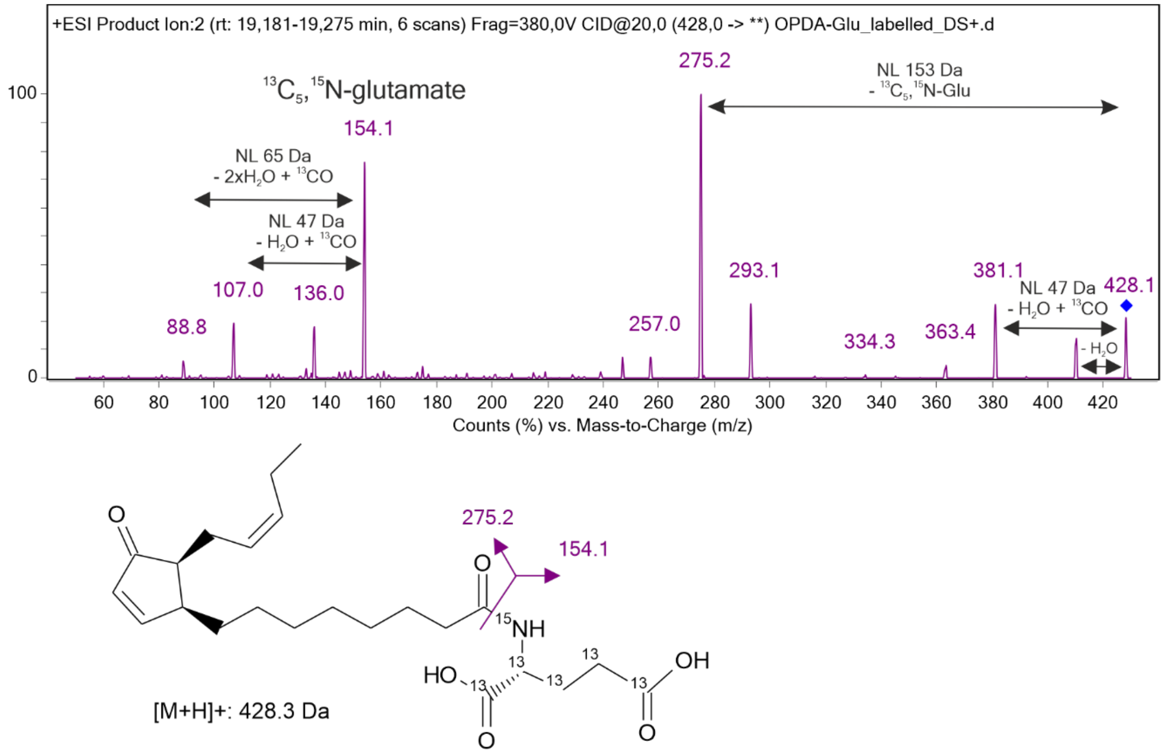


**Supplemental Note Figure 7** Fragmentation spectra of *cis*-(±)-OPDA-[$^{13}C_5$,$^{15}N$]*L*-glutamate at CE 20 eV.

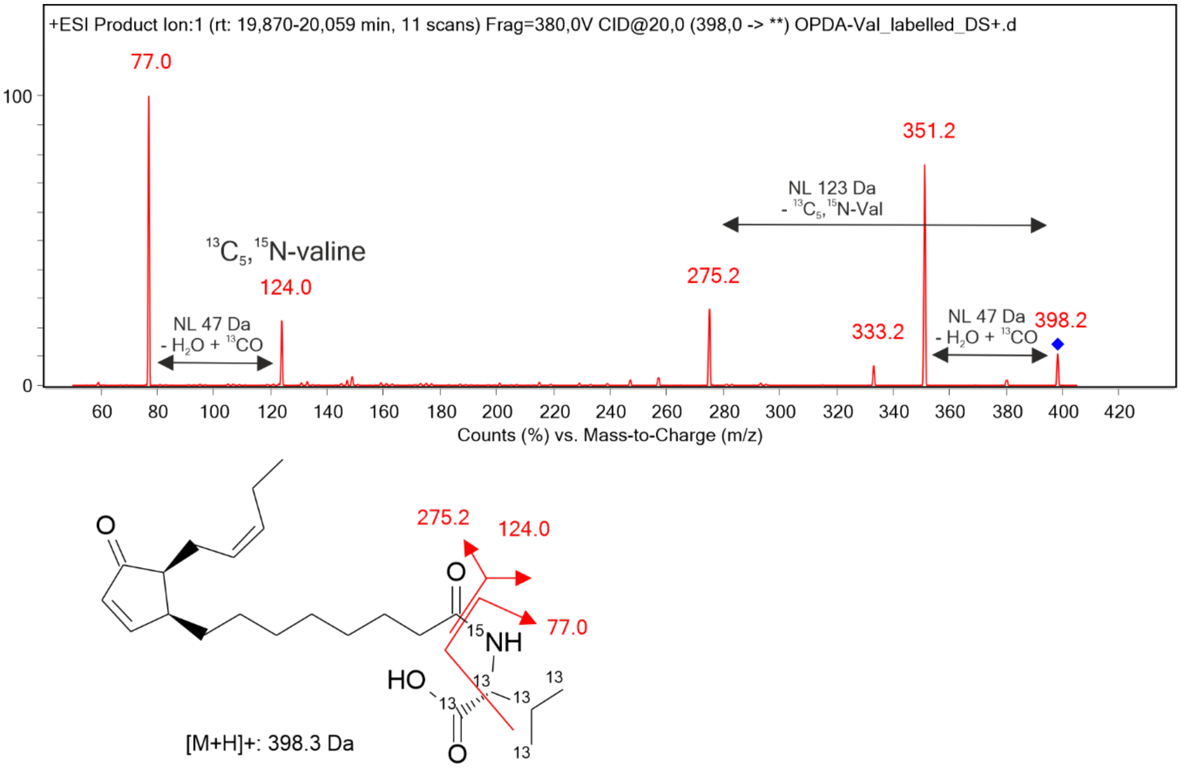


**Supplemental Note Figure 8** Fragmentation spectra of *cis*-(±)-OPDA-[$^{13}C_5$,$^{15}N$]*L*-valine at CE 20 eV.
