## Supplemental Data for "Conjugation of *cis*-OPDA with amino acids is a conserved pathway affecting *cis*-OPDA homeostasis upon stress responses"

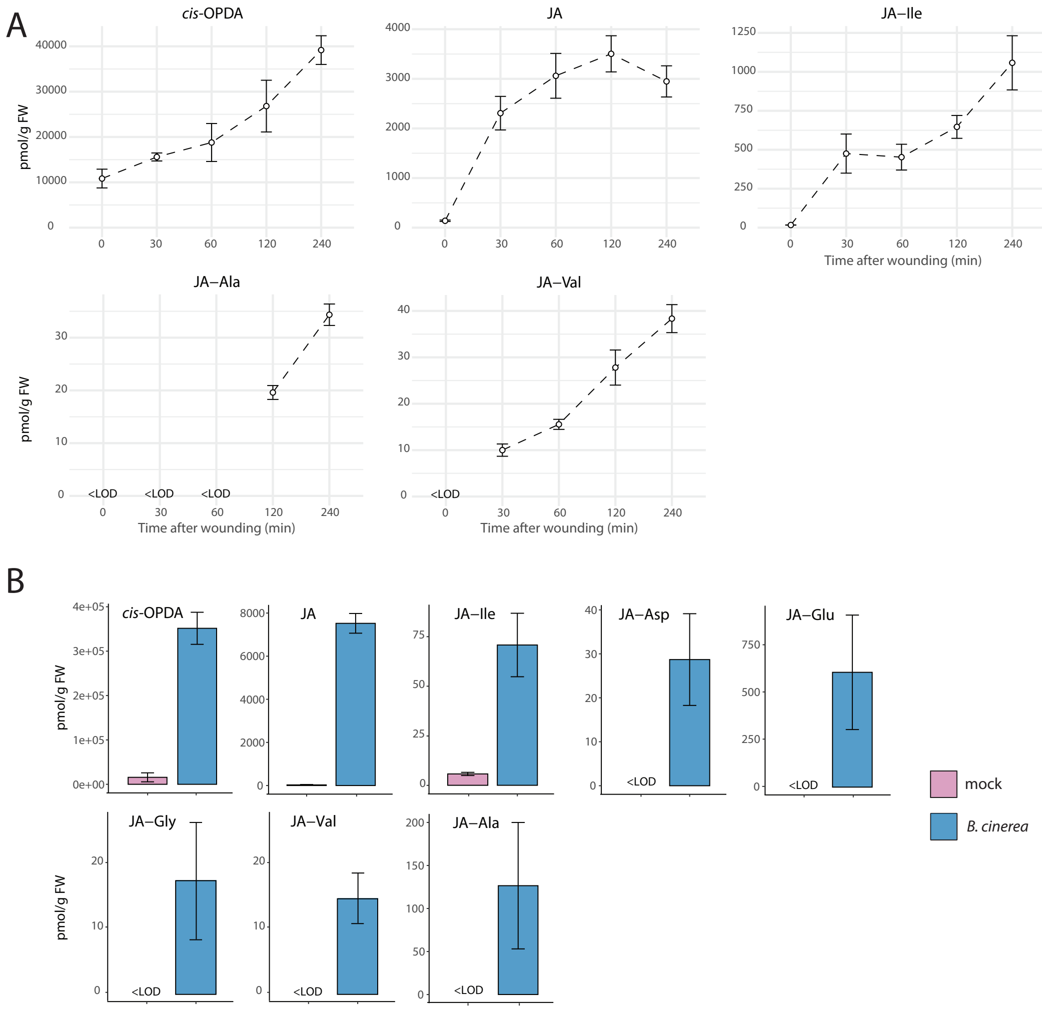


**Supplemental Figure S1** Accumulation of *cis*-OPDA, JA and JA-aa in Arabidopsis plants during stress responses. A, Time-course accumulation of *cis*-OPDA, JA, and indicated JA-aa in Col-0 after leaf wounding. Six-week-old plants were wounded, and damaged leaves were collected after the indicated times. B, Accumulation of *cis*-OPDA, JA, and indicated JA-aa in Col-0 plants infected with *B. cinerea* 3 days after inoculation.
Metabolite levels are expressed as pmoles per gram fresh weight (FW). Mean ± SD (*n*=3). Below the limit of detection, <LOD.

A

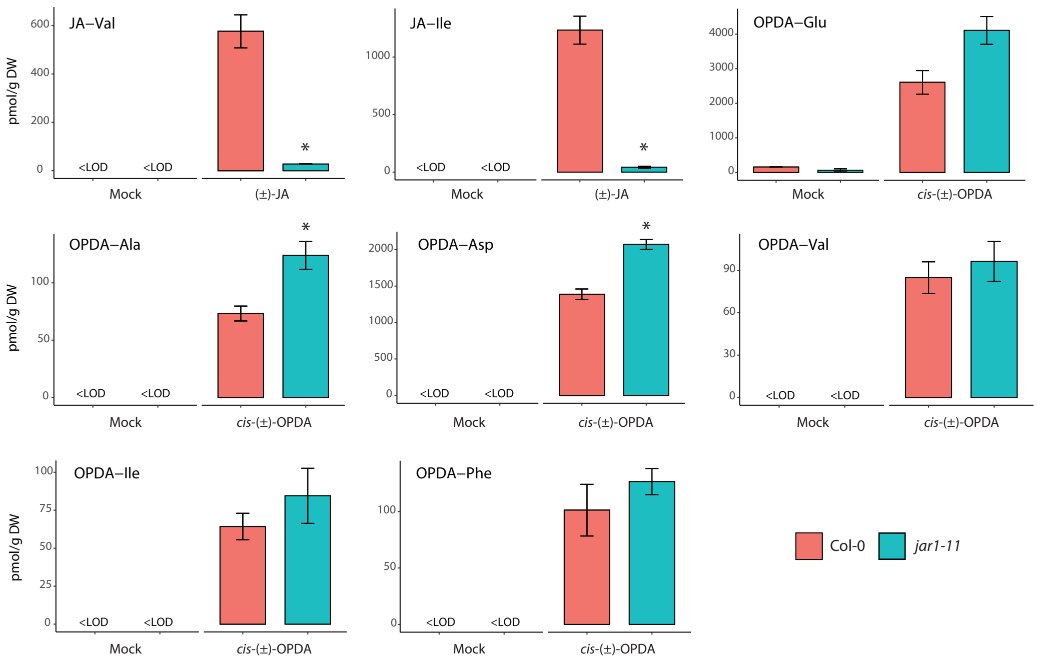


**Supplemental Figure S2** Amino acid conjugation of *cis*-OPDA is not mediated by JAR1/GH3.11 *in planta*. Accumulation of indicated JA-aa and OPDA-aa after exogenous treatment with or without 50 µM (±)-JA and *cis*-(±)-OPDA in *jar1-11* mutant. Asterisk indicates statistically significant differences, as determined by Student's *t*-test (Col-0 vs *jar1-11*; $P<0.05$). Metabolite concentrations are given as pmoles per gram dry weight (DW). Mean ± SD ($n$=3). Below the limit of detection, <LOD.

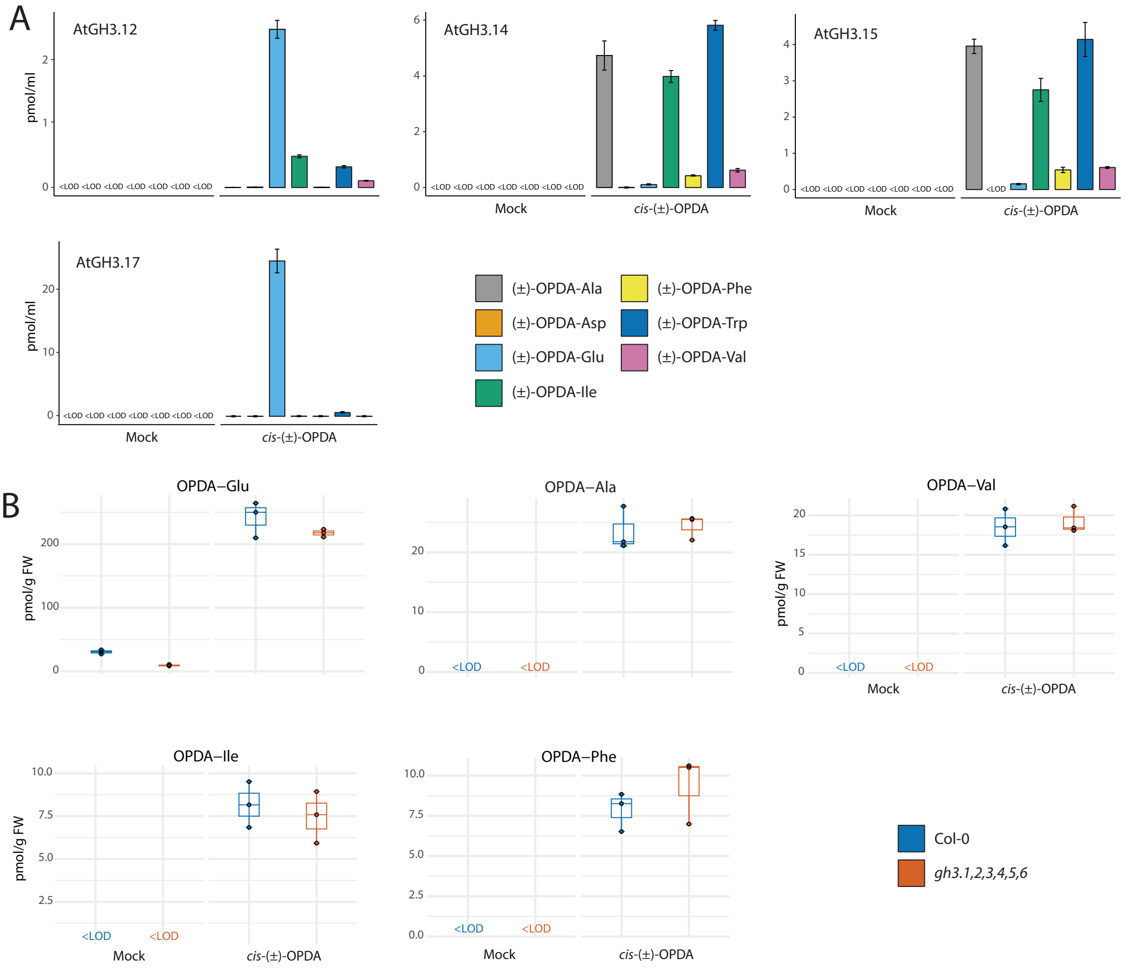


**Supplemental Figure S3** Conjugating activity with *cis*-(±)-OPDA of recombinant AtGH3s and accumulation of OPDA-aa in *gh3* sextuple mutant upon *cis*-(±)-OPDA feeding. A, Analysis of OPDA-aa synthesized by recombinant AtGH3.12, AtGH3.14, AtGH3.15, and AtGH3.17 in the bacterial assay. The cell lysate was incubated with or without 0.1 mM *cis*-(±)-OPDA and GH3 cofactor mixture for 5 h at 30 °C. The bacterial assay carried out with cell lysate from GFP-producing bacteria was used as a negative control. Cell lysate without *cis*-(±)-OPDA and cofactor mixture was used as a mock sample. OPDA-aa level is expressed as pmol/ml. B, Formation of OPDA-Glu, OPDA-Ala, OPDA-Val, OPDA-Ile, and OPDA-Phe after feeding of 7-day-old Arabidopsis Col-0 and *gh3* sextuple mutant (*gh3.1,gh3.2,gh3.3,gh3.4,gh3.5,gh3.6*) with or without 50 µM *cis*-(±)-OPDA for 3 h. OPDA-aa concentration is given as pmoles per gram fresh weight (FW). Horizontal lines in the box plots are medians, boxes show the upper and lower quartiles, and whiskers show the full data range.
Mean ± SD (*n*=3). Below the limit of detection, <LOD.

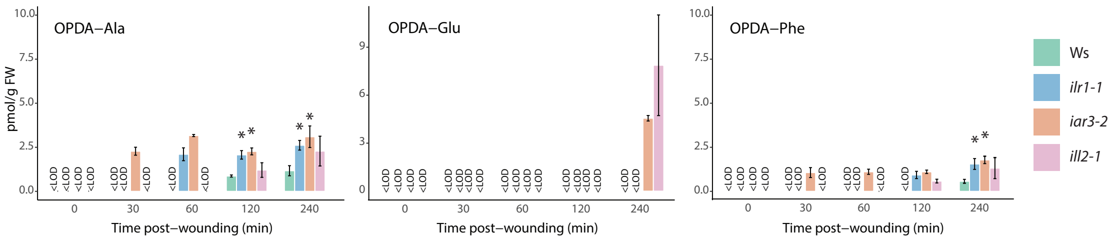


**Supplemental Figure S4** Accumulation of OPDA-aa in *ilr/ill* single knockout mutants upon wounding. Time-course accumulation of indicated OPDA-aa in Ws, *ilr-1*, *iar3-2*, and *ill2-1* single knockout mutants after leaf wounding. Six-week-old plants were wounded, and damaged leaves were collected after the indicated times. Asterisk indicates statistically significant differences, as determined by Student's *t*-test (Ws vs *ilr1-1*, *iar3-2,* or *ill2-1*; $P$<0.05). OPDA-aa concentrations are given as pmoles per gram fresh weight (FW). Mean ± SD ($n$=3). Below the limit of detection, <LOD.

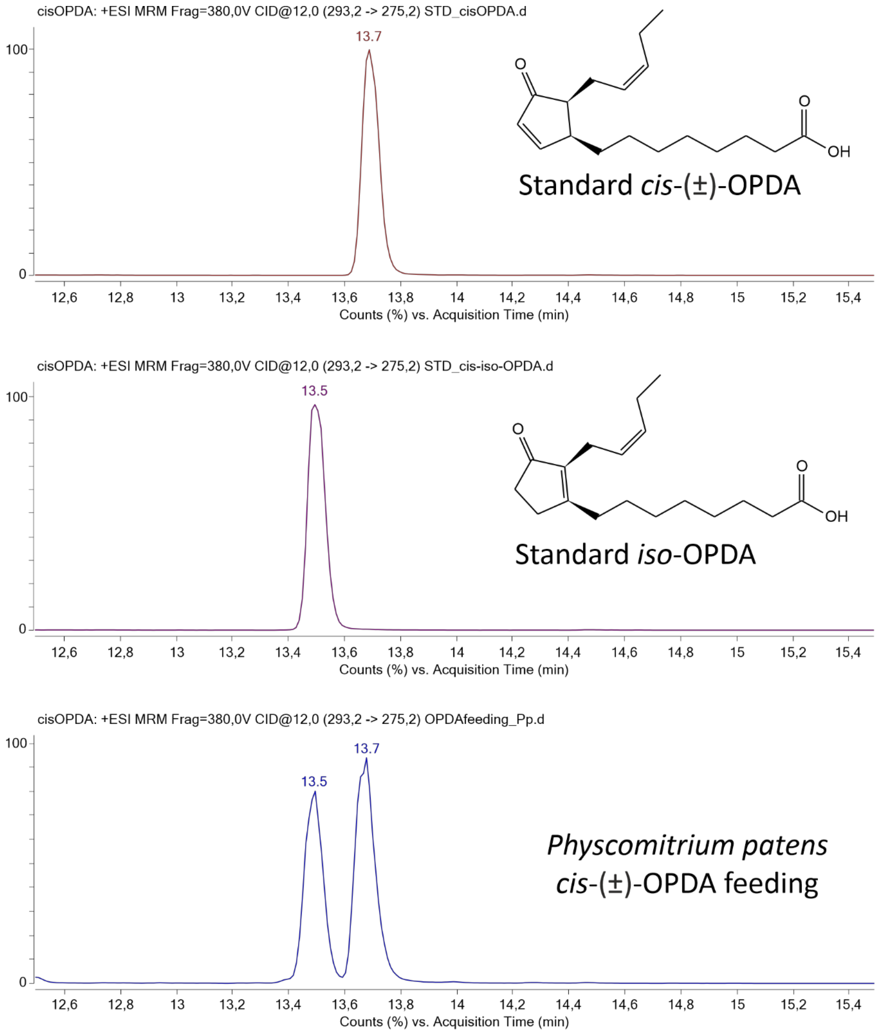


**Supplemental Figure S5** Extracted ion chromatogram of *cis-*/*iso*-OPDA and their identification in *Physcomitrium patens* after feeding with *cis*-(±)-OPDA (50 µM, 24 h) wild-type gametophores.

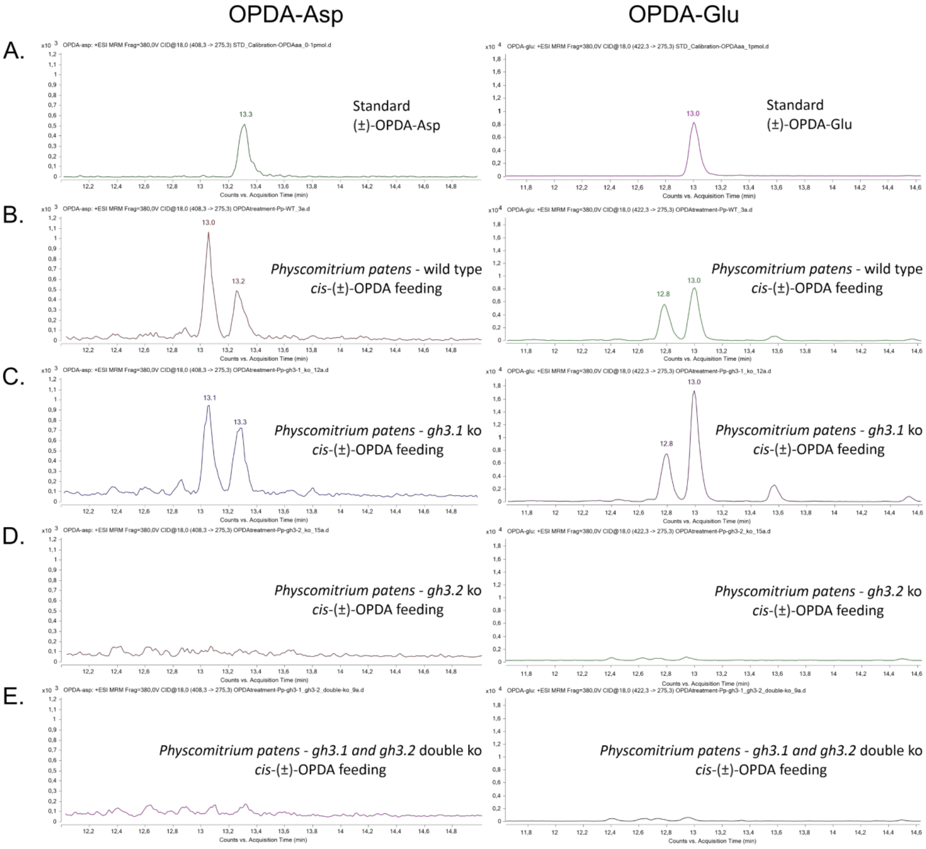


**Supplemental Figure S6** Extracted ion chromatogram of OPDA-Asp and OPDA-Glu standard (A) and *Physcomitrium patens* wild-type (B), *gh3.1* single (C), *gh3.2* single (D) and *gh3* double (E) knockout mutant gametophores after feeding with *cis*-(±)-OPDA (50 µM, 24 h).

**Supplemental Table S1** Semi-quantitative comparison of enzymatic activities with (±)-JA and amino acids of recombinant *Arabidopsis thaliana, Picea abies* and *Physcomitrium patens* GH3 proteins resulting from the bacterial assay.

| Species | GH3 | Group | Σ(+)-JA-Gly/ (-)-JA-Gly | Σ(+)-JA-Glu/ (-)-JA-Glu | Σ(+)-JA-Asp/ (-)-JA-Asp | (+)-JA-Ala | (-)-JA-Ala | (+)-JA-Val | (-)-JA-Val | (+)-JA-Met | (-)-JA-Met | Σ(+)-JA-Trp/ (-)-JA-Trp | (+)-JA-Ile | (-)-JA-Ile | (+)-JA-Phe | (-)-JA-Phe |
|---|---|---|---|---|---|---|---|---|---|---|---|---|---|---|---|---|
| Arabidopsis | AtGH3.1 | II | – | – | – | – | – | – | – | – | – | – | – | – | – | – |
| | AtGH3.2 | II | – | – | + | – | – | – | – | + | – | – | – | – | – | – |
| | AtGH3.3 | II | – | – | + | – | – | – | – | – | – | – | – | – | – | – |
| | AtGH3.4 | II | – | – | – | – | – | – | – | – | – | – | – | – | – | – |
| | AtGH3.5 | II | – | – | – | – | – | – | – | – | – | – | – | – | – | – |
| | AtGH3.6 | II | – | – | – | – | – | – | – | – | – | – | – | – | – | – |
| | AtGH3.7 | III | + | + | – | ++ | – | + | – | + | – | ++ | + | – | – | – |
| | AtGH3.8 | III | – | – | – | – | – | – | – | – | – | – | – | – | – | – |
| | AtGH3.9 | II | – | – | – | – | – | – | – | – | – | – | – | – | – | – |
| | AtGH3.10 | I | – | – | – | – | – | + | + | + | + | – | +++ | +++ | – | – |
| | AtGH3.11 | I | – | – | – | + | + | ++ | ++ | +++ | + + | ++ | ++++ | ++++ | +++ | + |
| | AtGH3.12 | III | – | ++ | – | – | – | + | + | + | + | – | + | + + | – | – |
| | AtGH3.13 | III | – | – | – | – | – | – | – | – | – | – | – | – | – | – |
| | AtGH3.14 | III | – | – | – | + | – | + | – | ++ | + | ++ | + | + | – | – |
| | AtGH3.15 | III | + | ++ | – | +++ | ++ | ++ | + | ++++ | +++ | ++++ | +++ | + | ++ | – |
| | AtGH3.16 | III | – | – | – | – | – | – | – | – | – | – | – | – | – | – |
| | AtGH3.17 | II | – | + | – | – | – | – | – | – | – | – | – | – | – | – |
| | AtGH3.19 | III | – | – | – | – | – | – | – | – | – | – | – | – | – | – |
| *P. abies* | PaGH3.16 | II | – | – | – | – | – | – | – | – | – | – | – | – | – | – |
| | PaGH3.17 | II | – | – | – | – | – | – | – | – | – | – | – | – | – | – |
| | PaGH3.gII.8 | II | – | – | – | – | – | – | – | – | – | – | – | – | – | – |
| | PaGH3.gII.9 | II | – | – | – | – | – | – | – | – | – | – | – | – | – | – |
| *P. patens* | PpGH3.1 | I | – | – | – | – | – | – | – | – | – | – | – | – | – | – |
| | PpGH3.2 | I | – | ++++ | ++ | + | + | + | + | + | + | ++ | + | + | – | – |
| Negative control | GFP | | – | – | – | – | – | – | – | – | – | – | – | – | – | – |

Range of detected (±)-JA-aa levels: < 0.1 pmol/mL (+), 0.1 – 1 pmol/mL (++), 1 – 10 pmol/mL (+++), > 10 pmol/mL (++++), and not detected (–).

**Supplemental Table S2** List of primers sequences used for qPCR analysis and cloning.

| Name | Gene ID | Purpose | Primer name | Sequence (5'–>3') |
|---|---|---|---|---|
| GRX480 | AT1G28480 | qPCR | qGRX480_Fw | TGATTGTGATTGGACGGAGA |
| | | | qGRX480_Rv | TAAACCGCCGGTAACTTCAC |
| ZAT10 | AT1G27730 | qPCR | qZAT10_Fw | ATCAACACTAGTAGCGTGTCC |
| | | | qZAT10_Rv | AGTCAAACCGAGGCTTCTTCG |
| THI2.1 | AT1G72260 | qPCR | qTHI2.1_Fw | GTTGGGTAAACGCCATTCTCG |
| | | | qTHI2.1_Rv | GTGGGACTACATAGCTCTTGG |
| PDF1.2 | AT5G44420 | qPCR | qPDF1.2_Fw | CACCCTTATCTTCGCTGCTC |
| | | | qPDF1.2_Rv | GTTGCATGATCCATGTTTGG |
| JAZ5 | AT1G17380 | qPCR | qJAZ5_Fw | CCAAGCCAGAGATTGTAACCG |
| | | | qJAZ5_Rv | CGAGATCTTTCGAACTTTGGC |
| VSP1 | AT5G24780 | qPCR | qVSP1_Fw | TCTCATACTCAAGCCAAACGG |
| | | | qVSP1_Rv | AGTATCCTCAACCAAATCAGC |
| ACT2 | AT3G18780 | qPCR | qAtACTIN2_Fw | GACCAGCTCTTCCATCGAGAA |
| | | | qAtACTIN2_Rv | CAAACGAGGGCTGGAACAAG |
| IAR3 | AT1G51760 | Cloning | IAR3_Δ25N_Fw | AACAGGTCTCACGCGTCCTCTAATGGGTTATCTCAA |
| | | | IAR3_Δ25N_Rv | AACAGGTCTCTATTATCAAAGTTCATCTTTTTTGTT |
| ILL6 | AT1G44350 | Cloning | ILL6_Δ25N_Fw | AACAGGTCTCACGCGACCAACTTACCTTTCTTTGAA |
| | | | ILL6_Δ25N_Rv | AACAGGTCTCTATTATTATGAATGTTTATCATTTAA |
| ILR1 | AT3G02875 | Cloning | ILR1_Δ25N_Fw | AACAGGTCTCACGCGTACGATTCTGGTTCGGGTCTC |
| | | | ILR1_Δ25N_Rv | AACAGGTCTCTATTACTATAATTCACTCTTAACCTCT |
| ILL2 | AT5G56660 | Cloning | AtILL2_Δ25N_Fw | AACAGGTCTCACGCGTGGATCGCCGAAGATACGTCT |
| | | | AtILL2_Δ25N_Rv | AACAGGTCTCTATTATTAGAGTTCTTCATGAAAGCC |
